# Functional dynamics of the eukaryotic RNA exosome upon complex formation

**DOI:** 10.64898/2026.10.10.758228

**Authors:** Jobst Liebau, Svetlana Kucher, Gloria Candy Danielle Sedoh, Enrica Bordignon, Remco Sprangers

**Affiliations:** Department of Biophysics I, Regensburg Center for Biochemistry, University of Regensburg, Universitätsstraße 31, 93053 Regensburg, Germany; Department of Biochemistry and Biophysics, Stockholm University, Svante Arrhenius väg 16C, 106 91 Stockholm, Sweden; Department of Physical Chemistry, University of Geneva, 30 Quai Ernest Ansermet, 1211 Geneva, Switzerland

**Keywords:** protein dynamics, RNA exosome, methyl-TROSY, CEST, relaxation dispersion, EPR

## Abstract

Rather than being static, proteins adopt multiple conformations and undergo dynamics on a wide range of timescales. Such conformational changes can be routinely probed by solution state NMR for protein sizes below 40 kDa and can be extended to large protein complexes with selective methyl-labeling strategies. Here, we expand the limit of methyl-NMR dynamics experiments to asymmetric protein complexes bigger than 300 kDa by investigating the inert core of the eukaryotic RNA exosome, Exo9. We demonstrate that isolated subunits are highly dynamic on the ms – s dynamics, which facilitates assembly. While ring subunits lose dynamics upon complex-formation, cap subunits and functional loops retain a large degree of flexibility as demonstrated by relaxation dispersion and chemical exchange saturation transfer experiments in the fully asymmetric Exo9 complex. Employing EPR spectroscopy we additionally revealed changes in the conformational ensemble and nanosecond dynamics of a spin-labeled functional loop region in the presence of the catalytic subunit Rrp44 and in response to substrate RNA binding. We thus demonstrate that an integrated NMR-EPR approach facilitates the characterization of functional dynamics that occur on a wide range of timescales in very large protein complexes.

## Main Text

The eukaryotic RNA exosome is involved in 3′ – 5′ mRNA degradation in the cytosol and in RNA processing and quality control in the nucleus (*1–3*). The structure of the exosome is well characterized for *S. cerevisiae*, *C. thermophilum* and *H. sapiens* orthologs in isolation or in combination with cofactors (*2*, *4–8*). It consists of an inert, nine-subunit core (Exo9), which is made up of six distinct subunits (Rrp41, Rrp45, Rrp43, Rrp46, Rrp42, Mtr3) that form a ring with a central channel that is stabilized by three distinct cap subunits (Csl4, Rrp4 and Rrp40) containing RNA-binding domains. Single-stranded RNA is passed via the cap subunits through the central exosomal channel and, at the other end, handed over to a tenth catalytic subunit, Rrp44, that hydrolyses RNA processively (**Fig. 1**) (*1*, *3*, *9*, *10*). In addition to the processive exonucleolytic activity, Rrp44 contains a PIN domain that conveys endonucleolytic activity and has recently been shown to hydrolyse circular RNA (*11*, *12*). While the exosome structure is relatively well known, functional dynamics of the exosome remain more elusive. We have previously demonstrated that the exosome undergoes extensive nanosecond dynamics (*13*) and that dynamics in a loop region of subunit Rrp42 (Rrp42-EL) (**Fig. 1b**), which is invisible in a static cryo-EM structure, are modified by Rrp44 and RNA substrate in a functionally relevant manner (*8*). Additionally, we previously observed multiple resonances for a number of residues in isolated subunits, indicative of slow dynamics and suggesting that other dynamics might be present in the exosome complex.

**Figure 1:**
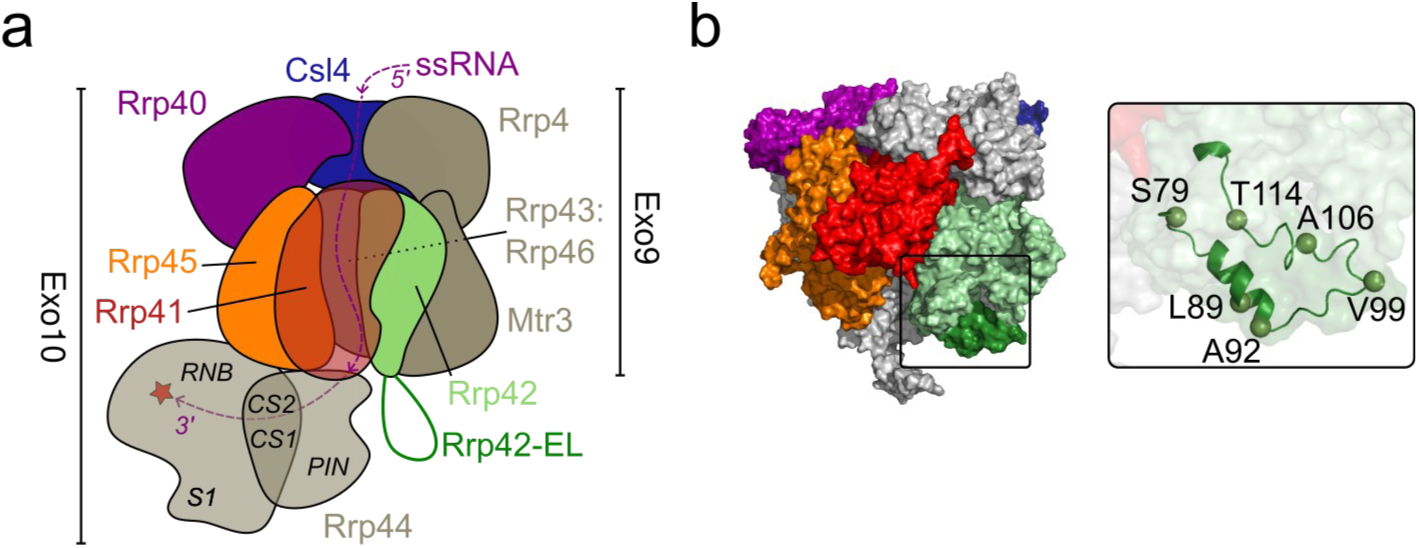
Static structure of the eukaryotic RNA exosome. **(a)** Schematic overview. Single-stranded RNA is passed via the cap subunits, through the exosomal channel formed by the ring subunits towards Rrp44, where it is degraded. Subunits studied here are shown in color (figure adapted from Ref. (*8*)). **(b)** Cryo-EM structure of *C. thermophilum* Exo9 (PDB: 8R1O). Invisible loops, such as Rrp42-EL (dark green), are modeled into the structure, as described in Ref. (*8*). Inset: Rrp42-EL (cartoon) in the open conformation. Residues that were mutated into Cys for EPR studies are shown as spheres.

Protein dynamics occur on a wide range of timescales, which are all crucial for protein function (*14–16*). For example, pico- to nanosecond dynamics have been employed as a proxy for the entropy of proteins (*17–19*) and are thus important determinants of binding, folding and misfolding with medical implications, e. g. for misfolded Aβ peptide that is involved in Alzheimer’s disease (*20*). Slower dynamics (µs – ms) are often relevant for the catalytic function of proteins as they constitute rate-limiting steps and often such dynamics correspond to collective motions of structural elements (*14*, *21*). Finally, proteins also exhibit slower, ms – s, dynamics, which can also be functionally relevant. For example, proteins can undergo folding transitions (*22*) or domain motions (*14*) and change folds (*23*) on that timescale. Furthermore, dynamics play a crucial role in ligand binding and complex assembly (*24*). Two distinct mechanisms can be distinguished to explain ligand binding or complex formation: in the conformational selection mechanism a protein constantly samples distinct, often sparsely populated states, which, however, show increased affinity to a substrate or binding partner. Binding then leads to selection and stabilization of the binding-competent state. In the induced-fit mechanism, a binding surface is present in the ground state of the protein and binding of substrate leads to a conformational rearrangement that stabilizes the bound state (*25–27*). Since NMR is specifically well suited to monitor sparsely populated states, it has been extensively used to analyze binding modes.

With the development of methyl-labeling methods in combination with methyl-TROSY experiments, NMR experiments have seen a substantial increase in sensitivity for proteins that are significantly bigger than the ∼40 kDa limit of ‘conventional’ NMR (*28–30*). Methyl-TROSY NMR facilitates the study of single-chained proteins as big as ∼100 kDa (*31*), highly symmetric protein complexes of 1 MDa size (*32*) and, as previously demonstrated by us, fully asymmetric protein complexes bigger than 400 kDa (*8*). A particular strength of NMR lies in the quantitative analysis of protein dynamics on a wide range of timescales, from picoseconds to hours, and recent method developments allowed quantification of dynamics also for large protein complexes. For example, pico- to nanosecond dynamics of methyl side chains can be probed by relaxation-violated coherence transfer experiments (*13*, *33*, *34*). Micro- to millisecond dynamics can be assayed by NMR using relaxation dispersion (RD) experiments. Various types of RD experiments have been developed, including single-quantum (SQ) Carr−Purcell−Meiboom−Gill (CPMG) (*35–37*), sensitivity-enhanced multi-quantum (MQ) CPMG (*38*) and spin-lock based R_1ρ_ RD experiments (*39*). Finally, NMR also allows to quantify ms – s dynamics by chemical exchange saturation transfer (CEST) or exchange spectroscopy (EXSY) experiments (*40*). Here, we focus on CEST experiments of methyl-labeled proteins, which provide two nuclei that can be monitored, ^1^H and ^13^C. CEST experiments for both nuclei have been developed (*41–43*) but require distinct labeling schemes when large protein complexes are investigated (^13^CHD_2_ labeling for ^13^C CEST, ^13^CH_3_ labeling for ^1^H CEST) and artifact suppression techniques in the case of ^1^H CEST (*42*). NMR experiments thus facilitate the study of a wide range of dynamics when dedicated experiments are combined with selective labeling schemes.

While we have previously demonstrated that pico- to nanosecond dynamics can be studied in large asymmetric complexes using relaxation-violated coherence selection experiments in methyl-labeled proteins (*13*), NMR is relatively insensitive to motions slower than a few nanoseconds and is blind in regions that do not contain methyl-bearing amino acids. Continuous wave (CW) electron paramagnetic resonance (EPR) spectroscopy at X band in combination with site-directed spin-labeling (SDSL) using nitroxides (*44*) is particularly well suited to investigate rotational dynamics in a time window from 30 ps to 30 ns (*45*) and thus complements NMR.

Here, we demonstrate that methyl dynamics experiments can be applied to quantify dynamics in individual subunits as well as the assembled RNA exosome core. We observe how dynamics are lost upon core exosome assembly, while motions in the periphery persist and are likely of functional relevance in the complex. Complementing the NMR data with site-specific information obtained by CW EPR, we provide functional insights into the dynamic properties of these large protein complexes.

## Results and Discussion

The formation of the exosomal complex was monitored by NMR using chemical shift perturbations as well as changes in dynamic properties for both isolated subunits and when reconstituted into the Exo9 complex. We first investigated the ring subunits Rrp41 and Rrp45, which in addition to being stable as monomers, can form a heterodimer. Next, we studied how complex formation affects the dynamic ensemble of the cap subunits Rrp40 and Csl4. Finally, we employ EPR to characterize functional dynamics in the Rrp42-EL loop that persist in the exosome in the absence and presence of the catalytic subunit and linear RNA substrate.

### Monomeric Rrp45 undergoes extensive millisecond dynamics

To probe interactions between Rrp45, Rrp41 and other Exo9 subunits we acquired methyl-TROSY spectra of Rrp45 as monomer, as heterodimer with Rrp41 and as protomer reconstituted into Exo9 (**Fig. 2a**). Upon dimer formation with Rrp41, Rrp45-I225 located at the binding interface shifts substantially and only a weak additional shift is seen in Exo9 suggesting that native contacts are formed in the dimer (**Fig. 2a**). Rrp45-I111, which is near the binding interface to Rrp41 also shifts upon dimer formation but shows additional shifts when Exo9 is formed. Other residues do not display strong perturbations upon dimer formation and native contacts are only formed in Exo9.

**Figure 2:**
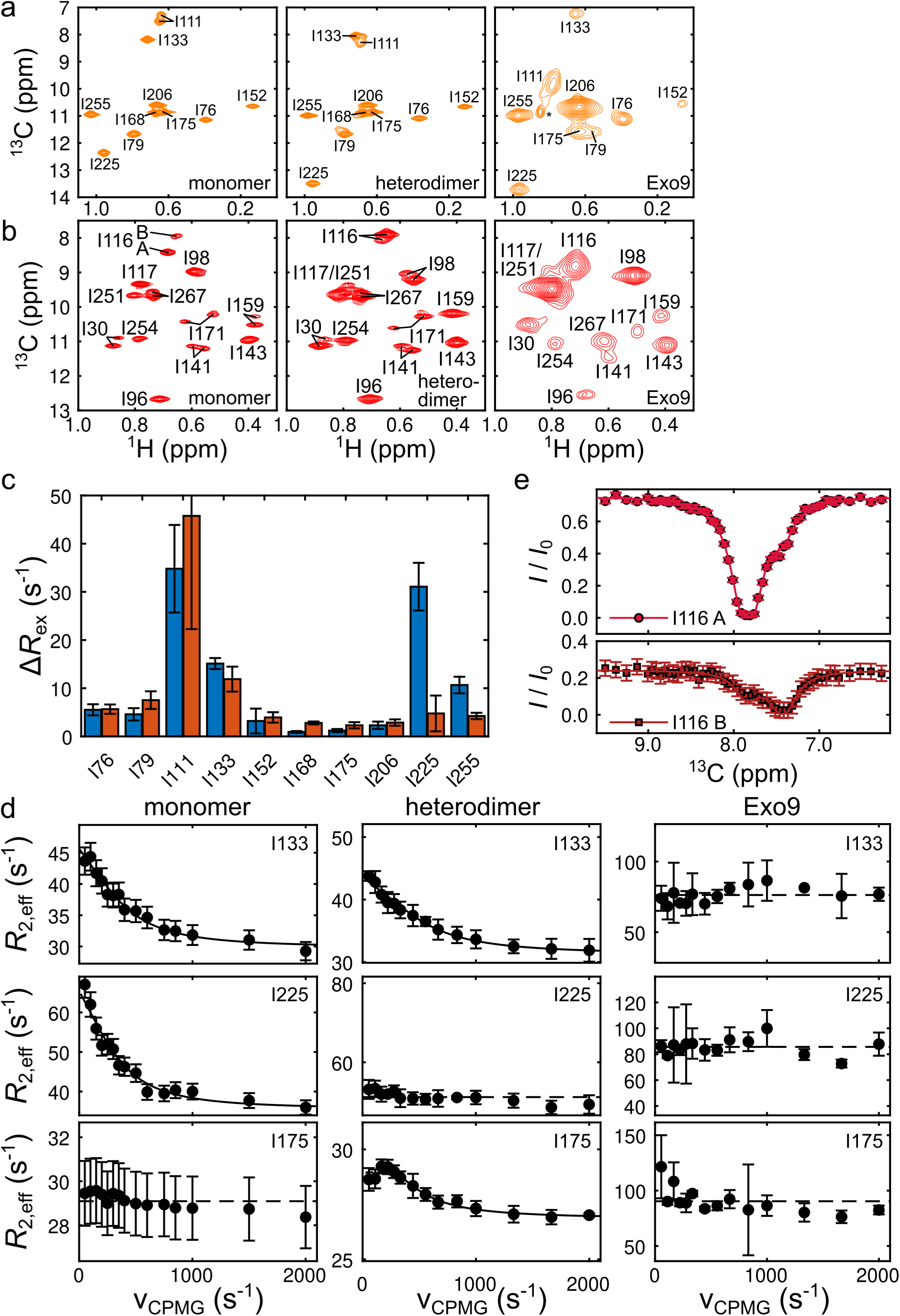
Dynamics of the ring subunits Rrp45 and Rrp41. **(a)** Methyl-TROSY spectra of I-labeled Rrp45 as monomer (left), heterodimer with Rrp41 (center) and reconstituted into Exo9 (right). * indicates an unassigned resonance. **(b)** Methyl-TROSY spectra of I-labeled Rrp41 as monomer (left), heterodimer with Rrp45 (center) and reconstituted into Exo9 (right). **(c)** Amplitude of the RD profiles, Δ*R*_ex_, in the Rrp45 monomer (blue) and heterodimer (red). **(d)** Representative SQ (monomer: column 1) and MQ (heterodimer: column 2, Exo9: column 3) RD profiles for I133 (row 1) that is dynamic in monomer and dimer, I225 (row 2) that shows no dynamics in the dimer and in Exo9 and I175 (row 3) that shows increased dynamics in the dimer compared to the monomer. Solid lines represent fits to a two-site exchange model (see *Methods*). Note that the asymptotic *R*_2_ rate at high ν_CPMG_ (*R*_2,inf_) depends on the protein size and type of experiment. Monomer and dimer spectra cover the same *R*_2eff_ range. **(e)** ^13^C CEST of the two conformations A and B of I116 in monomeric Rrp41 as labeled in **b**, acquired with *B*_1_ = 10 Hz. The fits are obtained for a two-site exchange model as described in the method section. Error bars are estimated from the signal-to-noise ratio of the spectra and represent ±1 SD.

In order to assess dynamics of the Rrp45 monomer, we conducted methyl RD experiments and grouped residues into two classes: one class of residues showing pronounced RD profiles with amplitudes ΔR_ex_ > 5 s^-1^ and one class showing no or weak profile amplitudes (**Fig. 2c**). Qualitatively, this analysis shows that parts of Rrp45 undergo millisecond dynamics between chemically distinct states, while other parts experience similar chemical environments. In order to quantify the exchange dynamics, we attempted to fit a global two-site exchange model to all the residues in the dynamic class, however, a single global fit did not converge. We then separated the dynamic residues into two groups, which provided convergent fits with similar exchange rates yet different populations of the excited state. This indicates that Rrp45 adopts two sparsely populated states (**Fig. 2d**, **SI Fig. S1, SI Table S5**) in addition to the ground state.

In addition to millisecond dynamics, monomeric Rrp45-I111 displays two resonances (**Fig. 2a**) suggesting the presence of one additional conformation. Simulation of CEST spectra suggest that exchange rates down to ∼7 s^-1^ should be observable (**SI Fig. S2**), however ^13^C CEST experiments do not indicate any distinct exchange peaks demonstrating that the dynamics are slower than ∼7 s^-1^ or absent. Notably, both states of I111 undergo millisecond dynamics that can be fit with the same exchange rate and minor population (**SI Fig. S1**). As a monomer, Rrp45 thus adopts two roughly equally populated conformations that sample at least two sparsely populated minor states on the millisecond timescale.

### Monomeric Rrp41 undergoes slow exchange dynamics

While monomeric Rrp45 undergoes pronounced millisecond dynamics, RD profiles for Rrp41 are flat indicating absence of such dynamics (**SI Fig. S3**). However, several residues display two resonances implying that Rrp41 adopts multiple conformations as well (**Fig. 2b**). Notably, one group of Ile residues (I171, I141 and I267) adopts two roughly equally populated states, while for another group (I30, I116 and I159) populations are unequally distributed between the two conformations. Employing ^13^C CEST experiments, we characterized kinetics between the two conformations sampled by residue I116 by fitting a two-site exchange model simultaneously to the CEST profile of each resonance and obtain an exchange rate of ∼3 s^-1^ and a major state population of ∼76 % (**Fig. 2e, SI Table S6**). Monomeric Rrp41 thus exists in various conformations in solution that undergo slow exchange.

### Complex formation alters the Rrp41/Rrp45 conformational ensemble

Upon dimer formation with Rrp41 the conformational ensemble of Rrp45 is drastically altered. Several residues now display flat or attenuated RD profiles (**Fig. 2d SI Fig. S4**), which indicates that dimer formation stabilizes Rrp45. Notably, for I225 the formation of native contacts in the heterodimer leads to loss of dynamics. Two residues remote from the Rrp45 binding surface (I168, I175) show increased flexibility (**Fig. 2d, SI Fig. S4**), while residues that adopt chemical shifts intermediate between monomer and Exo9 (I111, I133) retain dynamics suggesting that dynamic properties are important for complex formation for residues that have not formed native interactions in the dimer. In Exo9 we do not observe any RD profiles for Rrp45 suggesting that dynamics are abolished upon native contact formation (**Fig. 2d, SI Fig. S5**).

Similar to Rrp45, the conformational ensemble of Rrp41 is affected by formation of the heterodimer. Residue Rrp41-I159 in the core of Rrp41 adopts a single population indicating a stabilization of Rrp41. The chemical environments of Rrp41-I116 become more similar suggesting that one of the states of Rrp41 pre-samples the Rrp45-bound conformation (**Fig. 2b**), which, however, is not the native interaction since chemical shifts of I116 in the dimer differ from Exo9. Other residues retain conformational equilibria in the dimer suggesting that native contacts are not formed (**Fig. 2b**). Notably, the residues that sample two conformations in the monomer also show substantial CSPs upon Exo9 formation suggesting that conformational dynamics are crucial for the adoption of the native state, while residues which only adopt one conformation in the monomer do not display substantial CSPs upon reconstitution into Exo9 (**Fig. 2b**). Upon Exo9 formation, all residues display one resonance only, i. e. only one conformation is retained.

As monomers, both Rrp41 and Rrp45 thus adopt a broad distribution of conformational states, yet exchange dynamics occur on vastly different timescales, where Rrp45 undergoes mostly millisecond dynamics, while Rrp41 dynamics are orders of magnitude slower. Upon dimerization the dynamic and conformational landscape is substantially reshaped, giving rise to conformational stabilization near the interaction surface, where native interactions are formed, while regions remote from the interaction surface retain conformational flexibility or even become more flexible, facilitating binding to neighbouring subunits. Upon formation of native contacts in Exo9, the conformational flexibility on the ms – s timescale is lost suggesting that the ring formed by 6 subunits in the exosome is rigid on that timescale.

### Cap subunits retain dynamics in Exo9

We next turned to investigations of the dynamic properties of the exosome cap. Methyl-TROSY spectra of I-labeled Rrp40 (**Fig. 3a**) undergo strong chemical shift changes upon reconstitution into Exo9 indicating that Exo9 forms extensive contacts with Rrp40. The chemical shift changes for reconstituted Csl4 compared to monomeric Csl4 (**Fig. 3e**) are less drastic than for Rrp40, nevertheless all domains of Csl4 interact with Exo9.

**Figure 3:**
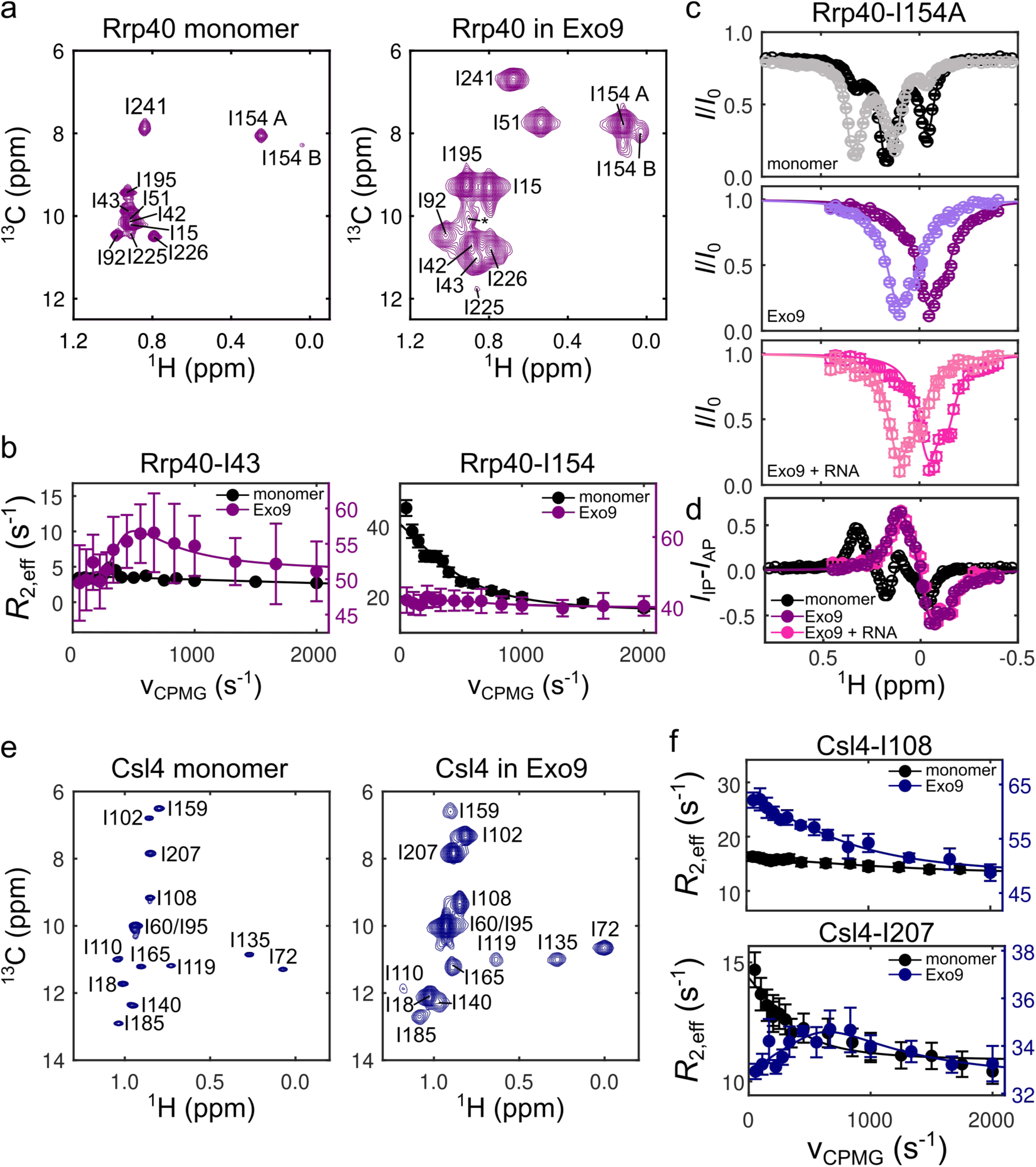
Dynamics of the cap subunits Rrp40 and Csl4. **(a)** Methyl-TROSY spectra of I-labeled Rrp40 as monomer (left) and reconstituted into Exo9 (right). * indicates unbound Rrp40. **(b)** RD profiles of selected residues in Rrp40 as monomer (black) and when reconstituted into Exo9 (purple). **(c)** ^1^H CEST acquired with *B*_1_ = 10 Hz for Rrp40-I154 as monomer (top) and when reconstituted into Exo9 in the absence (center) or presence (bottom) of RNA. AP profiles are in light color and IP profiles are in darker color. **(d)** ^1^H CEST IP/AP difference profiles of the data in **c**. Fits are to two-site exchange models as described in the method section. **(e)** Methyl-TROSY spectra of I-labeled Csl4 as monomer (left) and when reconstituted into Exo9 (right). **(f)** RD profiles of selected residues in Csl4 as monomers (black) and when reconstituted into Exo9 (blue).

Similar to monomeric Rrp41 and Rrp45, the cap subunits Rrp40 and Csl4 are highly dynamic as monomers. Both Rrp40 and Csl4 display pervasive millisecond dynamics as evidenced by RD experiments (**Fig. 3b, f, SI Fig. S6, S7**). Rrp40 consists of three domains, an N-terminal domain (NTD), an S1 and a KH domain, the latter two of which are RNA binding domains. While Ile residues in the NTD do not display millisecond motions, the RNA binding domains show millisecond motions in the Rrp40 monomer. In order to quantify dynamics, we conducted a global analysis (one *k_ex_*, *p_B_* for all residues), in which we disregarded the overlapping resonances. RD profiles of the non-overlapping resonances can be fit with a global exchange rate *k*_ex_ = 2600 s^-1^ and excited state population *p*_B_ = 41 %, suggesting that Rrp40 samples conformations in millisecond exchange (**SI Fig. S6, SI Table S7**). In addition to millisecond dynamics, Rrp40-I154 also undergoes slow exchange dynamics, which we probed by ^1^H CEST experiments (**Fig. 3c, d, Fig. S7**). Of note, in the ^1^H CEST experiments we observe experimental artifacts that arise for the anti-phase (AP) CEST profile at the position of the in-phase (IP) resonance and vice versa, which can likely be attributed to incomplete separation of the in-phase and anti-phase magnetization pathways during the experiment. Previous studies show less pronounced yet similar artifacts (*42*). We quantified the slow exchange using a two-site exchange model, in which we accounted for the incomplete magnetization separation employing a heuristic leakage factor (see *Methods*), and we obtained an exchange rate of ∼100 Hz and a minor population of ∼20 % (**Fig. 3c, SI Table S8**). The observation that Rrp40 samples conformations on the slow and intermediate-fast NMR timescale demonstrates the highly dynamic nature of the monomer.

Monomeric Csl4 undergoes millisecond dynamics as evidence by non-flat RD profiles for most of the Ile residues. However, a single global fit did not converge and we grouped the resonances into two classes, one of which, class A, undergoes very fast dynamics with a small minor population on the limit of detectability (*k*_ex_ = 8400 Hz and a minor population of ∼0.6 %) and another class B that undergoes slower dynamics with a higher second population (*k*_ex_ = 2300 Hz and a second population of ∼30 %) (**SI Fig. S8, SI Table S9**). Residues that belong to class A are located in the core of the domains of Csl4, which might indicate that Csl4 undergoes fast unfolding dynamics with a low unfolded population.

In contrast to the ring subunits, the cap subunits show some degree of stabilization when reconstituted into Exo9 but, at the same time, retain flexibility for some residues as evidenced by multi-quantum (MQ) RD experiments (**Fig. 3b, f, SI Fig. S9**). For Csl4, several residues, which are located in the S1 RNA binding domain (I108, I207) or N-terminal domain (NTD) (I18), that face the ring subunits of Exo9 undergo millisecond dynamics in Exo9.

Similar to Csl4, Rrp40 retains millisecond dynamics in Exo9, however, MQ RD profiles are less pronounced indicating that chemical environments of the major and minor populations are less distinct than for Csl4 in Exo9 or that the minor population is small (**SI Fig. S10**). Most notably, similar to Csl4, I43 in the Rrp40 NTD undergoes millisecond dynamics.

Notably, in the methyl-TROSY spectra of reconstituted Rrp40, we observe a minor population of monomeric Rrp40. ^1^H CEST spectra indicate that this minor conformation is in chemical exchange with Exo9-bound Rrp40 (**SI Fig. S11**). Currently, we cannot determine whether this observation is an experimental artifact or of biological relevance, however, it underlines the dynamic nature of Rrp40 binding to the exosome as we did not observe any residual unbound populations for other subunits after reconstitution. It further suggests that the millisecond dynamics observed for Rrp40 in Exo9 do not entail full detachment of Rrp40 from Exo9 since detachment dynamics are slower.

Rrp40-I154 retains slow exchange dynamics in Exo9, which we were able to demonstrate by ^1^H CEST experiments (**Fig. 3c, d and SI Fig. S7, SI Table S8**). To our knowledge this is the largest asymmetric complex, for which such dynamics have been observed by CEST experiments, demonstrating that NMR dynamics methods are applicable to large systems. Of note, the slow dynamics of Rrp40-I154 in Exo9 do not represent a detachment of the entire subunit either since chemical shifts of the exchanging conformations of Rrp40-I154 in Exo9 differ from those in the monomer. In the presence of substrate RNA, slow exchange dynamics of Rrp40-I154 in Exo9 persist (**Fig. 3c, d, SI Fig. S7, SI Table S8**). We currently cannot attribute a function to the conformational exchanges that occur in the cap subunits, yet we note that they are highly dynamic on the ms – s timescale, which contrasts with the rigid ring subunits and might play a role in substrate or cofactor recognition. Since dynamic Ile residues are located at interaction surfaces between Rrp40 and ring subunits, interface binding is dynamic and not rigid and may entail transient domain, but not full, detachment of Rrp40 and Csl4 on the millisecond timescale.

### Nanosecond Rrp42 loop dynamics depend on the catalytic subunit Rrp44

We previously demonstrated that one loop in Rrp42 (Rrp42-EL) exists in an equilibrium between closed and open conformations with respect to the Exo9 core (*8*). In the presence of Rrp44, the closed state becomes more populated; in the presence of RNA, Rrp42-EL only adopts the open state. This conformational equilibrium is of functional importance since Rrp42-EL blocks an aberrant access path to the catalytic subunit Rrp44, while it allows on-pathway RNA that takes the through channel route to access Rrp44. While the loop is not observable in our cryo-EM and X-ray crystallography structures, AlphaFold2 models suggest that it contains secondary structure elements, which, however, seem to unfold during a 100 ns MD simulation, as previously reported (*8*). Previously we also showed that ms – s dynamics play a crucial role for the function of the loop and paramagnetic relaxation enhancement data indicated that nanosecond dynamics, measured in terms of an order parameter, are altered in the closed state. However, it remained difficult to directly probe nanosecond motions of the loop by NMR. In order to characterize these dynamics, we employed electron paramagnetic resonance spectroscopy (EPR), which is sensitive to rotational motion on the nanosecond timescale.

To characterize the dynamics in Rrp42-EL, we introduced single Cys mutations into six positions in the loop in a Cys-free Rrp42^C59S^ construct (Rrp42^ΔC^) and labeled them with a proxyl radical (**Fig. 1b**). We then obtained CW EPR spectra for all the constructs in the monomeric variant proving the successful labeling **(SI Table S1)** and when reconstituted either into Exo9 or Exo10 in the presence or absence of RNA.

Among the studied singly-labeled monomeric Rrp42 constructs (**SI fig. 12**), the two labeled residues at both ends of the loop, S79C^PROXYL^ and T114C^PROXYL^ display more anisotropic spectra compared to the other positions located in the central loop, indicating restricted motions, hence slower dynamics, at both ends of the loop. When reconstituted into Exo9 or Exo10 (**Fig. 4a, SI fig. 13**) S79C^PROXYL^ and T114C^PROXYL^ display spectra characteristic for highly anisotropic slow motion independent on whether or not RNA is present. In contrast, the four sites L89C^PROXYL^, A92C^PROXYL^, V99C^PROXYL^ and A106C^PROXYL^ in the central region of the loop (**Fig. 4b, SI Fig. S13**) display a distinct major narrow-line spectral component (named ‘mobile’, with rotational correlation times of ∼1 ns, see **Fig. 4c**) and an additional component characterized by slower rotational dynamics (named ‘immobile’, with τ_C,immob_ ≈ 4-5 ns, see **Fig. 4c**), which are in a slow exchange (≳100ns) (**Fig. 4b, SI Fig. S13**). In Exo9, the spectra of the four labeled Rrp42 positions remain similar to the isolated proteins independent on the addition of RNA. In contrast, in Exo10 the spectra have a pronounced immobile component, which is suppressed in presence of RNA, giving a rise to a spectrum similar to Exo9 (**Fig. 4b inset, SI Fig. S13 insets**). These observations are reproduced on all four positions in the central loop region of Rrp42-EL (**SI Fig. S13**). A 2-component fitting of the spectra in Exo10 reveals that upon addition of RNA, the immobile component decreases by ∼23 % at 294 K (**Fig. 4b, d, SI Fig. 14**). This is in good agreement with previous NMR observations that showed that at 310 K 26 % of Rrp42-EL occupy the closed state in Exo10, while upon substrate addition only the open conformation persists. The residual immobile spectral fraction present in Exo10 after RNA addition, which is also visible in Exo9, might be attributed either to the transient formation of secondary structure elements as predicted by AlphaFold and MD, or to a bimodal rotameric population of the proxyl label side chain (*46*).

**Figure 4:**
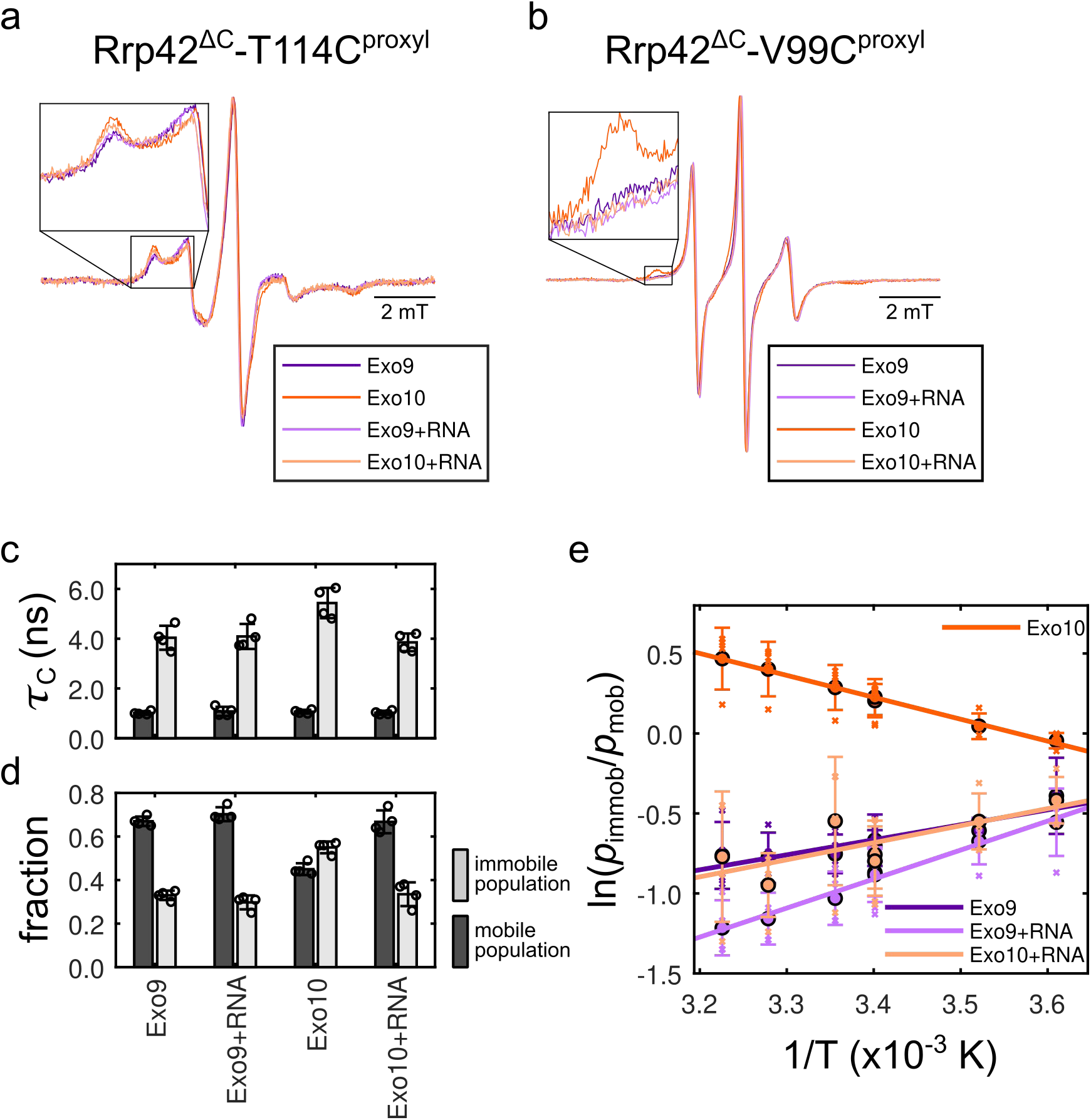
Nanosecond dynamics of Rrp42-EL. **(a)** Normalized EPR spectra of Rrp42^ΔC^-T114C^proxyl^, located at the edge of Rrp42-EL, in Exo9 (purple) and Exo10 (orange) with (dark colors) and without (light colors) RNA. Inset: Zoom into the spectral region of the immobile fraction. **(b)** EPR spectra of Rrp42^ΔC^-V99C^proxyl^, located in the center of Rrp42-EL, in Exo9 and Exo10 with and without RNA. Inset: Zoom into the spectral region of the immobile fraction. Color scheme as in **a**. **(c)** Correlation times and **(d)** fractions of the mobile (dark grey) and immobile (light grey) EPR spectral components obtained from spectral fits for the four probes in the central loop region of Rrp42-EL (L89C^PROXYL^, A92C^PROXYL^, V99C^PROXYL^ and A106C^PROXYL^) for spectra acquired at 294 K. Bar plots represent the mean of the parameters obtained from individual fits and error bars represent ±1 SD. **(e)** Van’t Hoff plot for the populations of the mobile (*p*_mob_) and immobile (*p*_immob_) fractions of Rrp42-EL in Exo9 and Exo10 with and without RNA. Circles are averages of measurements for four different proxyl positions in the loop (crosses represent the individual measurements). The corresponding rotational correlation times of mobile and immobile fractions are shown in an Arrhenius plot in **SI Fig. S14**. Error bars represent ±1 SD and fits are based on the van’t Hoff equation. Color scheme as in **a**.

To obtain insights into the thermodynamic properties of the equilibrium between the mobile and immobile populations of the loop, we detected CW EPR experiments at different temperatures (277-310 K) and analyzed the fractions of the two components in a van’t Hoff plot (**Fig. 4e, SI Table S10**). In Exo10 the transition from the mobile to the immobile loop is entropically driven by the catalytic subunit Rrp44. In the presence of RNA, interactions between the loop and Rrp44 are removed, as shown previously. Here, we observe that addition of RNA to Exo10 changes the free energy of the system and disfavors formation of the immobile loop. This situation is thermodynamically very similar to Exo9, i. e. RNA binding to Exo10 resets the thermodynamic equilibrium of Rrp42-EL back to the situation encountered in Exo9 and in Exo9 with RNA.

## Discussion

Dynamics on a wide range of timescales are essential for the function of enzymes and NMR has substantially contributed to demonstrate that proteins undergo functional dynamics on a broad range of timescales, however, systematic assessment of dynamics still remains scarce in particular in very large (>100 kDa) systems. Here, we demonstrate that a broad range of NMR dynamics experiments can be applied to large, asymmetric complexes. All subunits of the exosome undergo extensive dynamics as monomers (**Fig. 5a**), and some of these motions are essential for complex-formation based on a conformational selection mechanism though we cannot exclude that induced fit also plays a role. While dynamics are thus of mechanistic importance, we observe here that the opposite is true as well. Having formed a complex (**Fig. 5b**) the core of the exosome rigidifies and thus loses ms – s dynamics. Notably, we previously demonstrated that the core is still not rigid as nanosecond dynamics persist (*13*) and these may be of functional relevance as well since they contribute to the entropic component of the free energy. Loss of dynamics is thus a characteristic feature of complex formation in the exosome suggesting that dynamics are relevant for correct assembly but the subunits are rigid within the core of the exosome complex in agreement with the notion that one property of enzymes is to provide structural scaffolds that position or guide substrates to the catalytic site for efficient catalysis (*47*).

**Figure 5:**
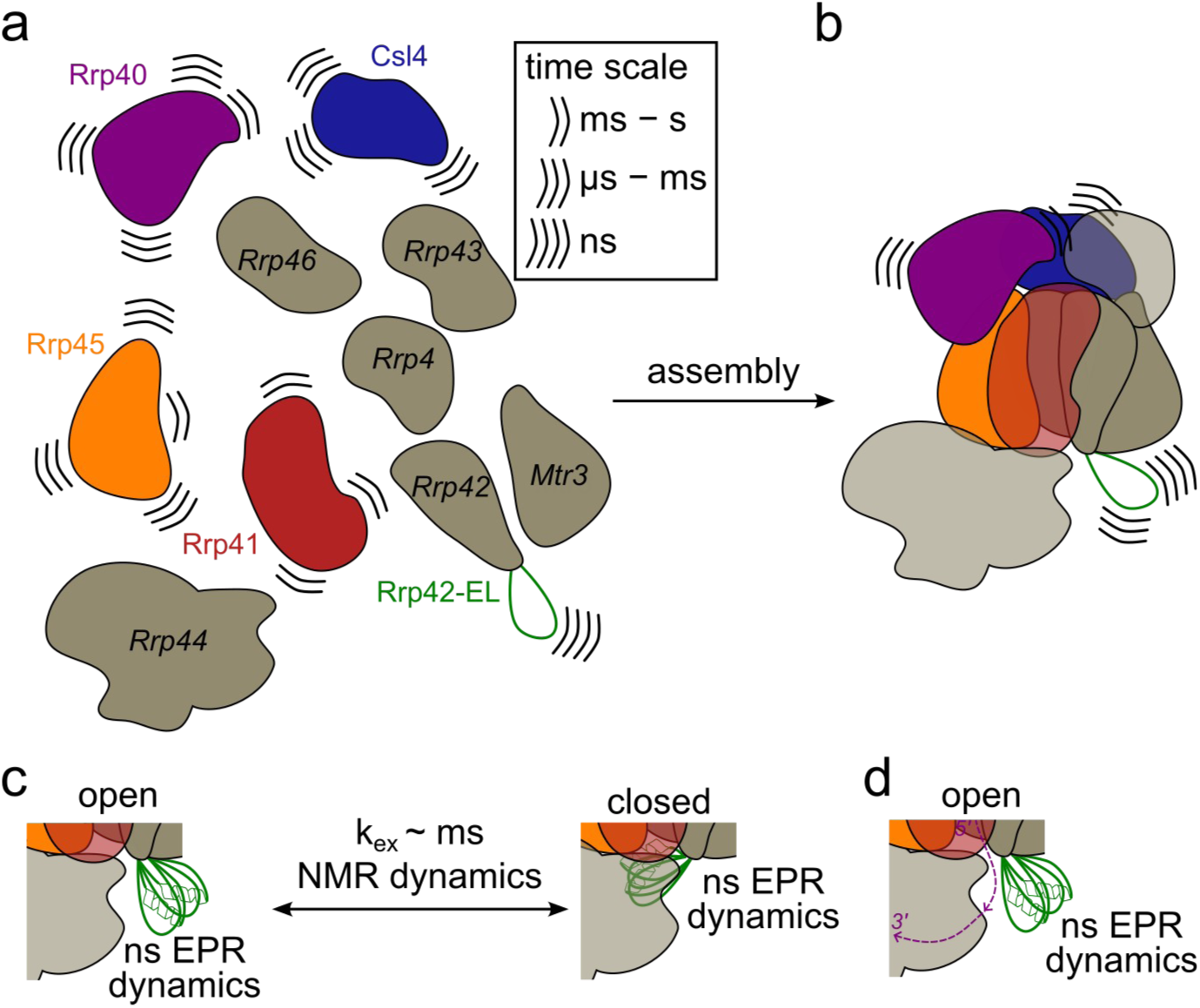
Exo9 rigidifies on assembly but retains a dynamic periphery. **(a)** In isolation, monomers of Exo9 are highly dynamic. Dynamics were investigated for the colored subunits/loops, while grey subunits were not studied here. **(b)** Upon assembly, the ring subunits rigidify but the periphery retains dynamics. **(c)** Rrp42-EL (green) retains extensive dynamics in the exosome complex undergoing exchange dynamics between an open and a closed state that can be quantified by NMR. In both conformations EPR-observable nanosecond dynamics persist, represented by multiple conformations of Rrp42-EL. **(d)** In the presence of single-stranded RNA (purple), Rrp42-EL adopts only the open conformation and retains EPR-observable dynamics.

In contrast to the exosome core, the periphery of the exosome, formed by the cap subunits and peripheral loop regions, retain extensive dynamics (**Fig. 5b**). Based on the cryo-EM structure (**Fig. 1b**), all domains of Rrp40 and Csl4 interact with ring subunits suggesting that this is the predominant conformation. However, our NMR dynamics experiments show that another minor conformation is sampled at the cap-ring interface. It has previously been observed in static cryo-EM structures that co-factor binding can lead to rearrangements of the cap subunits, thus possibly contributing to opening up of binding interfaces (*4*, *5*). For example, a substantial conformational change in a Csl4 loop upon Ski7 binding has been observed, suggesting that cap flexibility contributes to the formation of cofactor-binding interfaces (*48*). While we currently have no information about the structural features of that sparse conformation, we hypothesize that individual domains of the cap subunits can transiently detach from the ring subunits, which might play a role in substrate recruitment or cofactor binding, consistent with a notion that dynamic properties of enzymes provide gating mechanisms and determine substrate or cofactor binding (*49*).

While the relevance of the cap subunit dynamics remains to be investigated, we have previously demonstrated that Rrp42-EL serves as a dynamic plug that prevents RNA from accessing the exoribonucleolytic site of Rrp44 on an aberrant pathway while allowing passage of on-pathway RNA through the exosome channel towards Rrp44, rendering slow exchange dynamics functionally essential (*8*). Here, we complemented the dynamics studies of Rrp42-EL with EPR experiments that indicate the presence of two conformations independent on whether or not substrate or Rrp44 are present, although Rrp44 in the absence of RNA alters the population equilibrium. On the other hand, the NMR data explored dynamics, which cannot be observed directly by EPR, that demonstrate that Rrp42-EL undergoes millisecond exchange that is altered by the catalytic subunit Rrp44 and abolished in the presence of RNA substrate. The combination of NMR and EPR, however, facilitates characterization of the complex dynamic ensemble adopted by Rrp42-EL, which undergoes motions on the nanosecond to millisecond timescale giving rise to the following model (**Fig. 5c, d**): Rrp42-EL is in slow exchange on the EPR timscale (i.e. *k*_ex_ > 100 ns) between a flexible loop conformation and a more rigid conformation, in which it might transiently adopt α-helical secondary structure. The rigid conformation preferentially interacts with Rrp44 explaining the EPR-observed increase in the rigid component in apo Exo10, which is consistent with the increase in population of the closed state by ∼25 % observed by NMR. RNA addition to Exo10 displaces Rrp42-EL away from the exosome core and the loop is now in the open conformation, as observed by NMR. However, Rrp42-EL retains a high degree of flexibility in the open conformation as evidenced by EPR, which likely facilitates a re-binding once RNA has been processed. This dynamic feature is required to facilitate the displacement of the loop by RNA when inserted through the barrel but the loop also blocks an aberrant RNA access path in the apo exosome. The combination of NMR and EPR experiments that are sensitive to different motional timescales thus provided a comprehensive picture of dynamic processes in functionally essential loop regions.

## Conclusions

Dynamics are pervasive in biology, yet, in contrast to structural properties, they remain poorly characterized and their relevance for biological processes is thus long from being understood. This is particularly true for large protein complexes, in which dynamics are specifically challenging to study. Both NMR and EPR are suitable for the study of conformational ensembles and dynamics, yet on different timescales. Employing those complementary techniques, we demonstrated that conformational dynamics on a wide range of timescales can be characterized in very large and fully asymmetric protein complexes. This provided insights into complex formation mechanisms of the RNA exosome, in which subunits become rigid upon complex formation, but at the same time functional dynamics persist. Our study thus demonstrates that dynamics are dominant features also in large protein complexes, as the one studied here, and that loss of dynamics, can have a defined functional role.

## Materials and Methods

### Cloning, expression, methyl- and proxyl-labeling and purification

Single Cys point mutations were incorporated into Cys-free Rrp42^S52C^ (Rrp41^ΔC^) by a quick change-like procedure. Primers employed for design of mutants are listed in (**SI Table S1**).

*Chaetomium thermophilum* exosome subunits (**SI Table S2**) were prepared as described previously (*8*). Briefly, subunits were expressed individually or, for Rrp43 and Rrp46, in pairs, in D_2_O M9 medium employing recycled D_2_O and ^1^H^12^C glucose as carbon source for non-labeled subunits. Ile-δ1[^13^CH_3_]-labeled (I-labeled) subunits were expressed in fresh D_2_O M9 medium employing ^2^H^12^C glucose and, de-pending on the required labeling scheme, 60 mg/L ^2^H^12^C ketobutyric acid-4-^13^CH_3_ or ^2^H^12^C ketobutyric acid-4-^13^CHD_2_ were added 1 h prior to induction with isopropyl-β-D-thiogalactopyranoside (IPTG). For the preparation of EPR samples, cells were cultured in LB medium.

Protein was expressed overnight, cells were harvested, resuspended in 50 mM sodium phosphate buffer, pH 7.4, 150 mM NaCl (500mM NaCl for Rrp44^D168N-D536N^), 10 mM imidazole (buffer A), lysed by addition of Triton X100 and lysozyme and by sonication and debris was removed by centrifugation. The supernatant was applied onto a Ni-NTA column, washed with buffer A and for Rrp44^D168N-D536N^ with buffer A with 1 M NaCl, and protein was eluted in buffer A supplemented with 300 mM imidazole. 1 mg of TEV protease was added to the sample, which was dialysed overnight into 20 mM HEPES, pH 7.5, 150 mM NaCl (buffer B). TEV protease and the cleaved His-tag were removed by an inverse Ni-NTA column and the protein was subjected to size-exclusion chromatography into 10 mM HEPES buffer pH 7.5, 200 mM NaCl (buffer C).

For EPR, purified Rrp42^ΔC^, for which Cys residues were introduced into various positions of Rrp42-EL, were spin-labeled prior to SEC by adding 3-maleimido-2,2,5,5-tetramethyl-1-pyrrolidinyloxy (3-maleimido-PROXYL) in 5 times excess over the protein concentration. The sample was incubated at room temperature under gentle agitation for 1 hour and excess label was removed by SEC.

### RNA *in vitro* transcription

46mer linear RNA (**SI Table S3**) was prepared by *in vitro* transcription and purified by anion exchange chromatography using a preparative DNAPac 100 column (Dionex) as described previously (*8*).

### Exosome reconstitution

To reconstitute Exo9, the individual subunits were pooled in stoichiometric amounts, incubated for 30 minutes and subjected to size-exclusion chromatography. The fractions containing the exosome complex were pooled and concentrated.

For EPR, RNA and/or catalytically inactive Rrp44^D168N-D536N^ was added in 1.5 and/or 1.2 times excess, respectively, and all constructs, including apo Exo9, were subjected to another SEC purification step in order to remove larger aggregates that do not affect NMR spectra but might impact EPR experiments. Peak fractions were pooled and concentrated to a final concentration of 30 – 60 µM.

For NMR experiments, the sample was diluted 75x in D_2_O-based buffer C. For RNA-containing samples, RNA was added in 1.5x excess over the exosome complex and the sample was incubated for 30 minutes. Next, the sample was concentrated and the dilution-concentration procedure was repeated once to obtain the final samples used for subsequent NMR measurements.

An overview of all experiments conducted in this study is listed in **SI Table S4A**.

### NMR spectrometers

NMR experiments were conducted on 600 MHz and 800 MHz Avance Neo spectrometers (14.1 T and 18.8 T magnetic field strength, respectively) equipped with triple resonance cryogenic TCI probeheads cooled with liquid helium (800 MHz) or liquid nitrogen (600 MHz). Experiments were acquired with Topspin 4.0.3 (800 MHz spectrometer) or Topspin 4.0.8 (600 MHz spectrometer).

NMR experiments were conducted in NMR tubes with a diameter of 3 mm or in 5 mm Shigemi tubes. The final NMR sample volume was ∼180 µl (3 mm tubes) or ∼260 µl (5 mm Shigemi tubes) containing 100 – 200 µM Exo9, in which one subunit (Csl4, Rrp41 or Rrp45) was I-labeled, in D_2_O-based buffer C with 1.5x excess 46mer RNA where applicable. Experiments were conducted at temperatures indicated in **SI Table S4A**.

### Methyl-TROSY NMR

2D methyl-TROSY spectra were collected using the SOFAST-HMQC pulse sequence (*50*) with carbon acquisition times of 30 ms (Exo9) or 60 ms (exosome monomers) and an interscan delay of 0.5 s.

### Relaxation dispersion and CEST experiments

Constant time Carr-Purcell-Meiboom-Gill (CPMG) relaxation dispersion (RD) experiments were conducted with a relaxation delay (*T*_CPMG_) of 40 ms (SQ RD experiments) or 36 ms (MQ RD experiments) using at least 14 frequencies (ν_CPMG_). The maximum frequency that was used was 2000 Hz and the minimum frequency depended on the length of *T*_CPMG_ (**SI Table S4B**). CPMG RD experiments were conducted on a 800 MHz spectrometer except for monomeric Csl4, for which experiments were conducted on a 600 MHz spectrometer.

For ^1^H chemical exchange saturation transfer (CEST) experiments a ^13^CH_3_ I-labeling scheme was implemented, while ^13^C CEST experiments were conducted on ^13^CHD_2_ I-labeled samples. CEST profiles were conducted at *B*1 field strengths as indicated in the figures/captions (10–25 Hz) applied for *t*CEST = 400 ms on an 800 MHz spectrometer. CEST intensities were referenced to the intensity determined for offsets at ±10,000 Hz.

### NMR data analysis

NMR data were processed using the NMRPipe/NMRDraw software suite (version 11.7) (*51*). Methyl resonance intensities were obtained with NMRPipe.

For model fitting, in-house Matlab scripts were employed. In all fitting routines, the target function

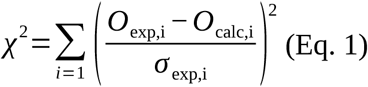

was minimized using the fminsearch routine in Matlab (version R2022b). In **Eq. 1** *O*_exp,i_ corresponds to an experimentally determined data point (*i*) in the recorded datasets *exp* {SQ/MQ RD data: *R*_2,eff_, ^1^H/^13^C CEST: normalized intensities}. *O*_calc,i_ corresponds to a back-calculated value of an observable based on the model parameters, as described below. *σ*_exp,i_ is an estimate of the measurement uncertainty for a data-point and is based on the noise level in the spectra or duplicate measurements.

A two-site exchange process between a ground state *G* and an excited state *E* as described by the equilibrium *G*⇄*E*, with *k*_ex_=*k*_EG_+*k*_GE_, 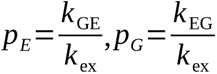, and *p_E_*=1 *− p_G_* was assumed in the fitted models.

### 1H and ^13^C CEST analysis

To fit the CEST data, the signal intensities (*I*) at offsets *ω*_CEST_ were back-calculated according to (*52*):

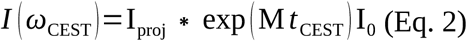

where *t*_CEST_ is the time during which the weak *B*_1_ field is applied. The equilibrium *z* magnetization is I =(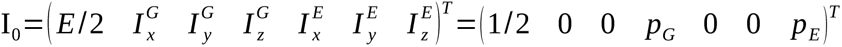, where *E* is the identity operator and 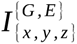 are the components of the magnetization vector in *x*, *y*, or *z* direction for the *G* and *E* states, respectively. M is the evolution matrix (according to the Bloch-McConnell equations):

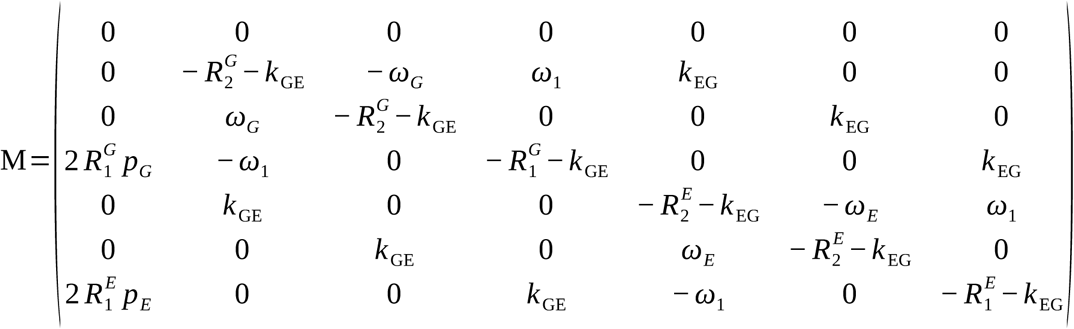

where, 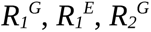 and 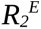 are the longitudinal and transverse relaxation rates of state *G* and *E*, respectively. *ω_G_*=*ω*_CEST_ *−ω_G_* and *ω_E_*=*ω*_CEST_ *−ω_E_* (in rad/s) denote the offsets between the CEST frequency and the chemical shifts of the ground and excited states, respectively. *ω*_1_=*γ*_C/H_ *B*_1_ is the weak *B*_1_ field (in rad/s) applied from the *y*-direction and *γ*_C/H_ is the gyromagnetic ratio of ^13^C or ^1^H. I_proj_ =(0 0 0 1 0 0 0) is the vector that projects the magnetization onto 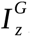, which results in observable magnetization.

For ^1^H CEST experiments we separately determined CEST profiles for in-phase (IP) and anti-phase (AP) magnetization, in order to suppress artifacts that may arise from NOE effects, as previously described (*52*). However, in the experimental implementation, we observe that the separation of IP and AP pathways is incomplete. In order to account for this leakage in the fitting procedure, we introduced a heuristic leakage factor λ and the observed intensities were calculated as:

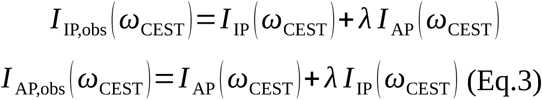

where *I*_AP_ and *I*_IP_ are calculated according to **Eq. 2**.

### Relaxation dispersion experiments

SQ and MQ CPMG relaxation dispersion (RD) data were back-calculated numerically using the Bloch-McConnell formalism, as described earlier (*53*). For global fits of SQ RD data of multiple resonances a single exchange rate, *k*_ex_, and excited state population *p*_E_ was assumed to exist, while Δω and *R*_2,inf_ were local parameters for each resonance. While MQ CPMG relaxation dispersion data was fit to a local two-site exchange model, the limited dataset size does not allow for accurate quantification and we therefore do not report any fit parameters for such experiments here but only qualitatively observe the existence or absence of dynamics.

### EPR experiments

Continuous wave (CW) EPR spectra were recorded on a Bruker ELEXSYS E500 X-Band Spectrometer equipped with a super high Q cavity ER 4122 SHQ. The spectra were acquired with a modulation amplitude of 1.5 G and a microwave (MW) power of 2 mW on samples of 20 μL volume filled into 0.9 mm glass capillaries. The temperature was set using a nitrogen flow cryostat and a temperature controller (Eurotherm). The Exo9/10 samples spectra were measured in a temperature range from 277 K to 310 K, and the integrity of the samples was confirmed by verifying the spectral reproducibility at 294 K after each temperature series.

The labeling efficiency was estimated as the ratio of spin concentration to protein concentration. The spin concentration was determined from double integration of the CW EPR spectra recorded at 294 K, with respect to a 4-Hydroxy-TEMPO (Sigma-Aldrich, CAS: 2226-96-2) standard of known concentration (100 μM) dissolved in water.

The spectra were fitted using 2 components with the chili function of Easyspin6.0.10 (*54*) in Matlab2025a. For the simulation, the following values of the g and A coupling tensors were fixed and employed for both components: *g_xx_* = 2.0075; *g_yy_* = 2.0055; *g_zz_* = 2.0028; *A_xx_* = 0.55 mT; *A_yy_* = 0.55 mT; *A_zz_* = 3.7 mT. The rotational correlation time and the weight of both components were varied and used subsequently for the van’t Hoff plot.

## Acknowledgments

We thank Nadine Stefan for excellent technical assistance.

## Authors contributions

Conceptualization: RS, JL

Methodology: RS, JL, SK, EB

Investigation: JL, SK, GS

Formal Analysis: JL, SK, GS

Visualization: JL, SK

Funding acquisition: RS, JL, EB

Project administration: RS, JL

Supervision: RS, JL

Writing – original draft: RS, JL, EB, SK

Writing – review & editing: RS, JL, EB, SK

## Data and materials availability

All data are available in the manuscript or the supplementary materials.

## Supplementary Materials

Supplementary Tables S1-S11

Supplementary Figures S1-S14

