## Supplementary Material for "Functional dynamics of the eukaryotic RNA exosome upon complex formation"

**Table of contents**

Supplementary Tables S1-S11 Page S2 – S13

Supplementary Figures S1-S14 Page S14 – S27

**Table S1: EPR mutants of Rrp42 with internal database reference, primers used for construct design, monomer concentrations and proxyl labeling efficiency.** See **Fig. S14** for monomeric EPR spectra.

| Construct | Internal database reference | Forward primer | Reverse primer | Protein concentration (µM) | Labeling efficiency (%) |
|---|---|---|---|---|---|
| Rrp42$^{\Delta C}$-S79C | #2835 | CGGAAGTGGAAAAAACCGTTTGCCGTTCGAAAGAAGATGAAG | CTTCATCTTCTTTCGAACGGCAAACGGTTTTTTCCACTTCCG | 112 | 108±10 |
| Rrp42$^{\Delta C}$-L89C | #2836 | GATGAAGAAGTCGGCTGCCTGGTTGCGAGCG | CGCTCGCAACCAGGCAGCCGACTTCTTCATC | 150 | 105±10 |
| Rrp42$^{\Delta C}$-A92C | #2837 | GTCGGCCTGCTGGTTTGTAGCGCCGGTGATATG | CATATCACCGGCGCTACAAACCAGCAGGCCGAC | 91 | 87±10 |
| Rrp42$^{\Delta C}$-V99C | #2838 | GAGCGCCGGTGATATGGACTGTGATGACGAAGAAG | CTTCTTCGTCATCACAGTCCATATCACCGGCGCTC | 117 | 95±10 |
| Rrp42$^{\Delta C}$-A106C | #2176 | GACGAAGAAGGCTATTGCAAAGTCGGTGCGGATAAC | GTTATCCGCACCGACTTTGCAATAGCCTTCTTCGTC | 100 | 103±10 |
| Rrp42$^{\Delta C}$-T114C | #2839 | GGTGCGGATAACCGTTGCGGCGAAGCATCATG | CATGATGCTTCGCCGCAACGGTTATCCGCACC | 117 | 75±10 |

**Table S2: Protein construct used in this study.**

| Protein | Accession code | Internal database reference |
|---|---|---|
| Rrp45 | G0S755 | #1215 |
| Rrp41 | G0SC21 | #1245 |
| Rrp43 | G0S1P1 | #1353 |
| Rrp46 | G0SCD1 | |
| Rrp42 | G0RZG4 | #1214 |
| Mtr3 | P0CT46 | #1246 |
| Rrp40 | G0RZX8 | #1213 |
| Rrp4 | G0S9A0 | #1233 |
| Csl4 | G0SE33 | #1247 |
| Rrp44 | G0SEX3 | #1242 |
| Rrp44$^{D168N, D536N}$ | | #1671 |

**Table S3: Sequence of RNA employed in this study.**

| RNA | Sequence | Internal database reference |
|---|---|---|
| 46mer | GGAGGAGAGGUGAGGAGAGAGGAGAGGAAGGAAGGGAAGAAAGAAG | #72 |

**Table S4: (A) Protein constructs and NMR/EPR experiments conducted on them in this study. (B) Parameters for relaxations dispersion experiments.**

**A**

| **Labeled construct** | **Labeling scheme** | **Non-labeled subunit(s) and substrate** | **Type(s) of experiment** |
|---|---|---|---|
| Rrp45 | $^{13}CH_3$ I-labeled | | Methyl-TROSY HMQC, SQ RD at 800 MHz, $^{1}H$ CEST at 298 K |
| Rrp45 | $^{13}CH_3$ I-labeled | Rrp41 | Methyl-TROSY HMQC, MQ RD at 800 MHz at 298 K |
| Rrp45 | $^{13}CH_3$ I-labeled | Exo9 | Methyl-TROSY HMQC, MQ RD at 800 MHz at 313 K |
| Rrp41 | $^{13}CH_3$ I-labeled | | Methyl-TROSY HMQC, SQ RD at 800 MHz, $^{1}H$ CEST |
| Rrp41 | $^{13}CHD_2$ I-labeled | | $^{13}C$ CEST at 298 K |
| Rrp41 | $^{13}CH_3$ I-labeled | Rrp45 | Methyl-TROSY HMQC, $^{1}H$ CEST at 298 K |
| Rrp41 | $^{13}CHD_2$ I-labeled | Rrp45 | $^{13}C$ CEST at 298 K |
| Rrp41 | $^{13}CH_3$ I-labeled | Exo9 | Methyl-TROSY HMQC at 313 K |
| Csl4 | $^{13}CH_3$ I-labeled | | Methyl-TROSY HMQC, SQ RD at 600 MHz at 313 K |
| Csl4 | $^{13}CH_3$ I-labeled | Exo9 | Methyl-TROSY HMQC, MQ RD at 800 MHz at 313 K |
| Rrp40 | $^{13}CH_3$ I-labeled | | Methyl-TROSY HMQC, SQ RD at 800 MHz, $^{1}H$ CEST at 298 K |
| Rrp40 | $^{13}CH_3$ I-labeled | Exo9 | Methyl-TROSY HMQC, MQ RD at 800 MHz, $^{1}H$ CEST at 313 K |
| Rrp40 | $^{13}CH_3$ I-labeled | Exo9 + RNA | Methyl-TROSY HMQC, $^{1}H$ CEST at 313 K |
| Rrp42$^{\Delta C\text{-}S79C}$ | proxyl | monomer, Exo9, Exo10, Exo9+RNA, Exo10+RNA | CW EPR at 277, 284, 294, 298, 305 and 310 K |
| Rrp42$^{\Delta C\text{-}L89C}$ | proxyl | monomer, Exo9, Exo10, Exo9+RNA, Exo10+RNA | CW EPR at 277, 284, 294, 298, 305 and 310 K |
| Rrp42$^{\Delta C\text{-}A92C}$ | proxyl | monomer, Exo9, Exo10, Exo9+RNA, Exo10+RNA | CW EPR at 277, 284, 294, 298, 305 and 310 K |
| Rrp42$^{\Delta C\text{-}V99C}$ | proxyl | monomer, Exo9, Exo10, Exo9+RNA, Exo10+RNA | CW EPR at 277, 284, 294, 298, 305 and 310 K |
| Rrp42$^{\Delta C\text{-}A106C}$ | proxyl | monomer, Exo9, Exo10, Exo9+RNA, Exo10+RNA | CW EPR at 277, 284, 294, 298, 305 and 310 K |
| Rrp42$^{\Delta C\text{-}T114C}$ | proxyl | monomer, Exo9, Exo10, Exo9+RNA, Exo10+RNA | CW EPR at 277, 284, 294, 298, 305 and 310 K |

**B**

| Labeled construct | Labeling scheme | unlabeled subunit(s) | Field (MHz) | Type | $\nu_{CPMG}$ | | Data points | $T_{CPMG}$ (ms) |
|---|---|---|---|---|---|---|---|---|
| | | | | | min (Hz) | max (Hz) | | |
| Rrp45 | $^{13}CH_3$ I-labeled | | 800 | SQ | 50 | 2000 | 15 | 40 |
| Rrp45 | $^{13}CH_3$ I-labeled | Rrp41 | 800 | MQ | 56 | 2000 | 14 | 36 |
| Rrp45 | $^{13}CH_3$ I-labeled | Exo9 | 800 | MQ | 56 | 2000 | 14 | 36 |
| Rrp41 | $^{13}CH_3$ I-labeled | | 800 | SQ | 50 | 2000 | 15 | 40 |
| Csl4 | $^{13}CH_3$ I-labeled | | 600 | SQ | 50 | 2000 | 17 | 40 |
| Csl4 | $^{13}CH_3$ I-labeled | Exo9 | 800 | MQ | 56 | 2000 | 14 | 36 |
| Rrp40 | $^{13}CH_3$ I-labeled | | 800 | SQ | 50 | 2000 | 15 | 40 |
| Rrp40 | $^{13}CH_3$ I-labeled | Exo9 | 800 | MQ | 56 | 2000 | 14 | 36 |

**Table S5: Global fit for relaxation dispersion data of Rrp45.** $k_{ex}$ and $p_B$ are global parameters for residues in the same group, $\Delta\nu_C$ and $R_{2,inf}$ are residues-specific parameters. I152, I168, I175 and I206 do not display any relaxation dispersion. See **Fig. S1** for RD profiles.

| **ID** | **Group** | **$k_{ex}$ ($s^{-1}$)** | **$p_B$** | **$R_{2,inf}$ ($s^{-1}$)** | **$\Delta\nu_C$ (ppm)** |
|---|---|---|---|---|---|
| 76 | A | 2449 | 0.10 | 31.5 | 0.31 |
| 79 | A | 2449 | 0.10 | 31.9 | 0.27 |
| 133 | A | 2449 | 0.10 | 29.8 | 0.53 |
| 255 | A | 2449 | 0.10 | 36.7 | 0.43 |
| 111 A | B | 2244 | 0.45 | 37.4 | 0.44 |
| 111 B | B | 2244 | 0.45 | 36.9 | 0.47 |
| 225 | B | 2244 | 0.45 | 35.6 | 0.41 |
| 152 | - | n.d. | n.d. | n.d. | n.d. |
| 168 | - | n.d. | n.d. | n.d. | n.d. |
| 175 | - | n.d. | n.d. | n.d. | n.d. |
| 206 | - | n.d. | n.d. | n.d. | n.d. |

**Table S6: $^{13}$C CEST fitting parameters for Rrp41-I116.**

| construct | $k_{ex}$ ($s^{-1}$) | $p_B$ | $\Delta\nu_C$ (ppm) | $R_2$ ($s^{-1}$) | $R_1$ ($s^{-1}$) |
|---|---|---|---|---|---|
| monomer | 3.4±0.4 | 0.242±0.004 | 2.57±0.01 | 44±4 | 0.3±0.3 |

**Table S7: Global fit for relaxation dispersion data of Rrp40.** $k_{ex}$ and $p_B$ are global parameters, $\Delta\nu_C$ and $R_{2,inf}$ are residues-specific parameters. I15, I42, I51 and I225 were not included in the fit due to peak overlap. I43 and I195 do not display any relaxation dispersion. See **Fig. S6** for RD profiles.

| **ID** | **Group** | **$k_{ex}$ ($s^{-1}$)** | **$p_B$** | **$R_{2,inf}$ ($s^{-1}$)** | **$\Delta\nu_C$ (ppm)** |
|---|---|---|---|---|---|
| 92 | A | 2569 | 0.41 | 22.1 | 0.15 |
| 154 A | A | 2569 | 0.41 | 16.3 | 0.42 |
| 154 B | A | 2569 | 0.41 | 15.5 | 0.38 |
| 226 | A | 2569 | 0.41 | 21.9 | 0.14 |
| 241 | A | 2569 | 0.41 | 49.4 | 0.28 |
| 15 | - | n.d. | n.d. | n.d. | n.d. |
| 42 | - | n.d. | n.d. | n.d. | n.d. |
| 43 | - | n.d. | n.d. | n.d. | n.d. |
| 51 | - | n.d. | n.d. | n.d. | n.d. |
| 225 | - | n.d. | n.d. | n.d. | n.d. |
| 195 | - | n.d. | n.d. | n.d. | n.d. |

**Table S8: Fit parameters for $^{1}$H CEST of Rrp40-I154 as monomer, in Exo9 and in Exo9 with RNA.** For the monomer, the major and minor state were fit globally and spin state leakage was accounted for by the empirical leakage parameter λ as lined out in the method section. See **Fig. S7** for CEST profiles.

| **construct** | **$k_{ex}$ ($s^{-1}$)** | **$p_B$** | **$\Delta\nu_C$ (ppm)** | **$R_2$ ($s^{-1}$)** | **$R_1$ ($s^{-1}$)** | **λ** |
|---|---|---|---|---|---|---|
| monomer | 113 | 0.19 | 0.21 | 25 | 1.1 | 0.28 |
| Exo9 | 19 | 0.23 | 0.08 | 222 | 3.9 | 0 |
| Exo9 + RNA | 133 | 0.10 | 0.12 | 91 | 3.3 | 0 |

**Table S9: Global fit for relaxation dispersion data of Csl4.** $k_{ex}$ and $p_B$ are global parameters for residues in the same group, $\Delta\nu_C$ and $R_{2,inf}$ are residues-specific parameters. I60 and I95 were not included in the fit due to peak overlap. I140 does not display any relaxation dispersion. See **Fig. S8** for RD profiles.

| ID | Group | $k_{ex}$ ($s^{-1}$) | $p_B$ | $R_{2,inf}$ ($s^{-1}$) | $\Delta\nu_C$ (ppm) |
|---|---|---|---|---|---|
| 102 | A | 2256 | 0.304 | 20.2 | 0.25 |
| 207 | A | 2256 | 0.304 | 10.9 | 0.25 |
| 18 | B | 8412 | 0.057 | 13.6 | 0.66 |
| 72 | B | 8412 | 0.057 | 21.4 | 0.90 |
| 108 | B | 8412 | 0.057 | 12.9 | 0.92 |
| 110 | B | 8412 | 0.057 | 19.1 | 0.85 |
| 119 | B | 8412 | 0.057 | 20.6 | 0.96 |
| 135 | B | 8412 | 0.057 | 21.7 | 0.75 |
| 159 | B | 8412 | 0.057 | 19.1 | 0.92 |
| 165 | B | 8412 | 0.057 | 18.9 | 0.85 |
| 185 | B | 8412 | 0.057 | 19.2 | 0.98 |
| 60 | - | n.d. | n.d. | n.d. | n.d. |
| 95 | - | n.d. | n.d. | n.d. | n.d. |
| 140 | - | n.d. | n.d. | n.d. | n.d. |

**Table S10: Van't Hoff fit parameters for Rrp42-EL in Exo9 and Exo10 in the presence and absence of RNA substrate. Free energy and the immobile loop population were calculated for 293 K.** Errors are given as ±1 SD.

| **Construct** | **ΔH (kJ mol$^{-1}$)** | **ΔS (J mol$^{-1}$ K$^{-1}$)** | **ΔG at 293 K (kJ mol$^{-1}$)** | **$p_{immobile}$ at 293 K** |
|---|---|---|---|---|
| Exo9 | -8±5 | -32±18 | 1.4 | 0.36 |
| Exo9 + RNA | -15±5 | -59±16 | 2.3 | 0.28 |
| Exo10 | 11±3 | 41±11 | -1.0 | 0.60 |
| Exo10 + RNA | -9±7 | -36±24 | 1.5 | 0.35 |

**Table S11: Arrhenius fit parameters for Rrp42-EL in Exo9 and Exo10 in the presence and absence of RNA substrate.** See **Fig. S14** for the Arrhenius plot.

| | **Immobile component** | | **Mobile component** | |
|---|---|---|---|---|
| **Construct** | **$E_a$ (kJ mol$^{-1}$)** | **ln(τ) (s)** | **$E_a$ (kJ mol$^{-1}$)** | **ln(τ) (s)** |
| Exo9 | 11±5 | -24±2 | 12±1 | -25.9±0.6 |
| Exo9 + RNA | 14±2 | -25.3±0.7 | 15±2 | -26.7±0.7 |
| Exo10 | 7±4 | -22±1 | 12±2 | -25.7±0.8 |
| Exo10 + RNA | 15±4 | -25±2 | 13±2 | -26.2±0.7 |

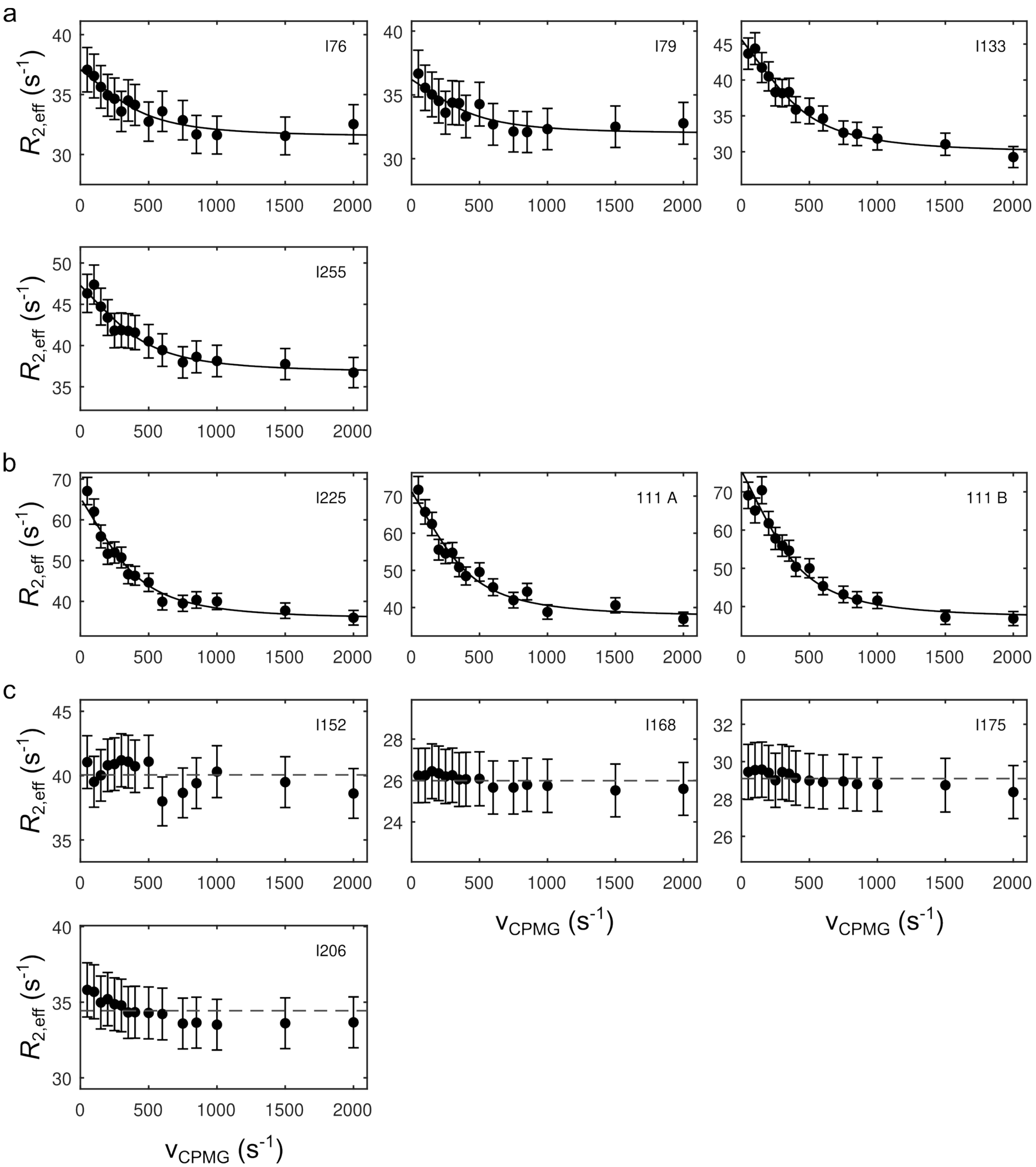


**Figure S1: Global fit for relaxation dispersion data of Rrp45. (a)** Resonances included in the global fit for **(a)** group A and **(b)** group B based on a two-state model as described in the *Method* section. **(c)** Resonances excluded from the global fit due to lack of relaxation dispersion. See **Table S5** for fitting parameters.

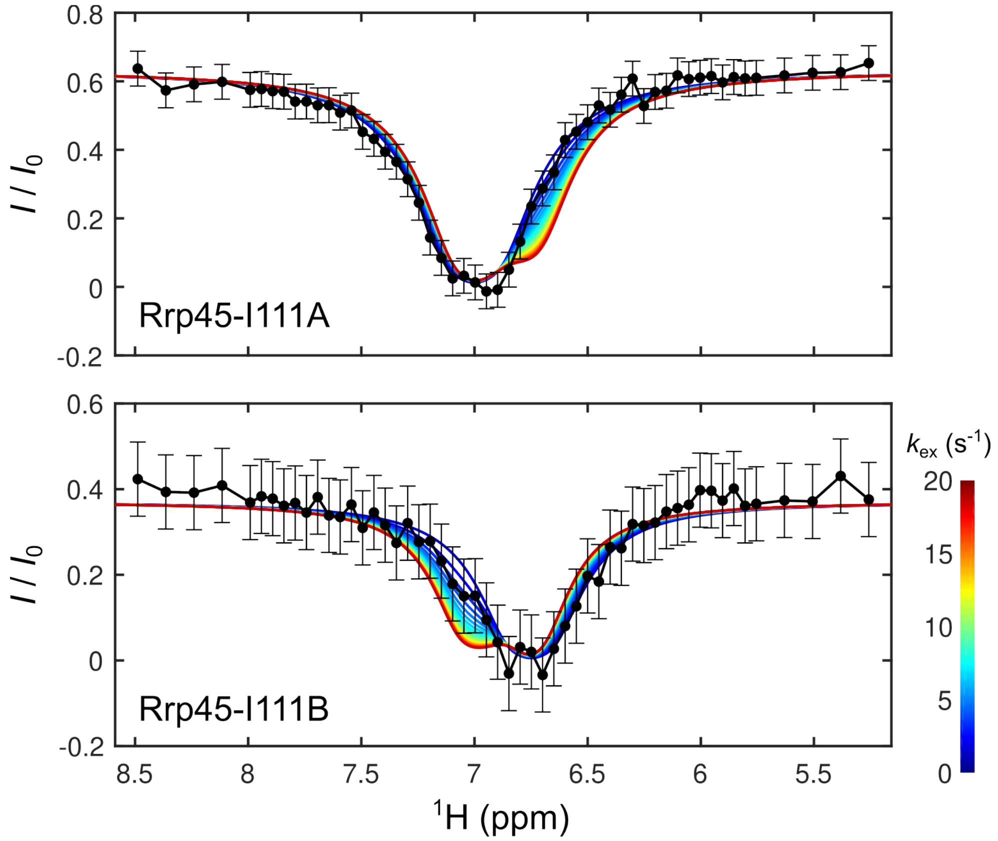


**Figure S2: Comparison of simulated (in color) and experimental (black) CEST data for Rrp45-I111 for $k_{ex}$ = 0 – 20 s$^{-1}$ (blue ⟶ red).** A model that assumes no exchange ($k_{ex}$ = 0, dark blue) can be fit well to the data and deviations between data and simulation are observed for $k_{ex} \gtrsim$ 7 s$^{-1}$ (light blue). Other simulation parameters were: $B_1$ = 10 Hz, $t_{CEST}$ = 0.4 s, $p_B$ = 0.37, $\Delta\nu_H$ = 0.25 ppm, $R_{1,A/B}$ = 1 s$^{-1}$, $R_{2A}$ = 80 s$^{-1}$, $R_{2B}$ = 60 s$^{-1}$.

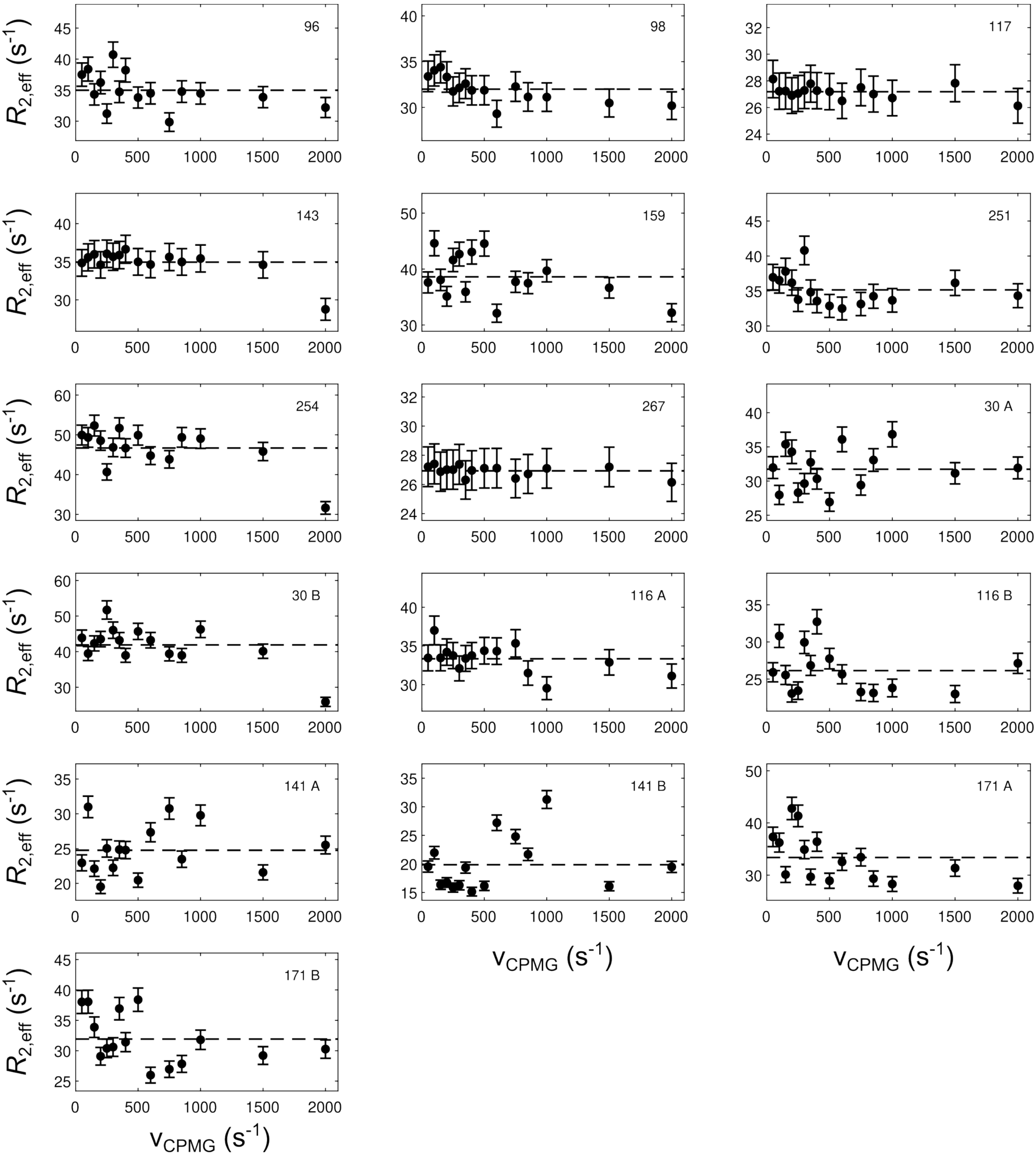


**Figure S3: SQ CPMG relaxation dispersion data for I-labeled monomeric Rrp41.** Dashed lines indicate absence of relaxation dispersion.

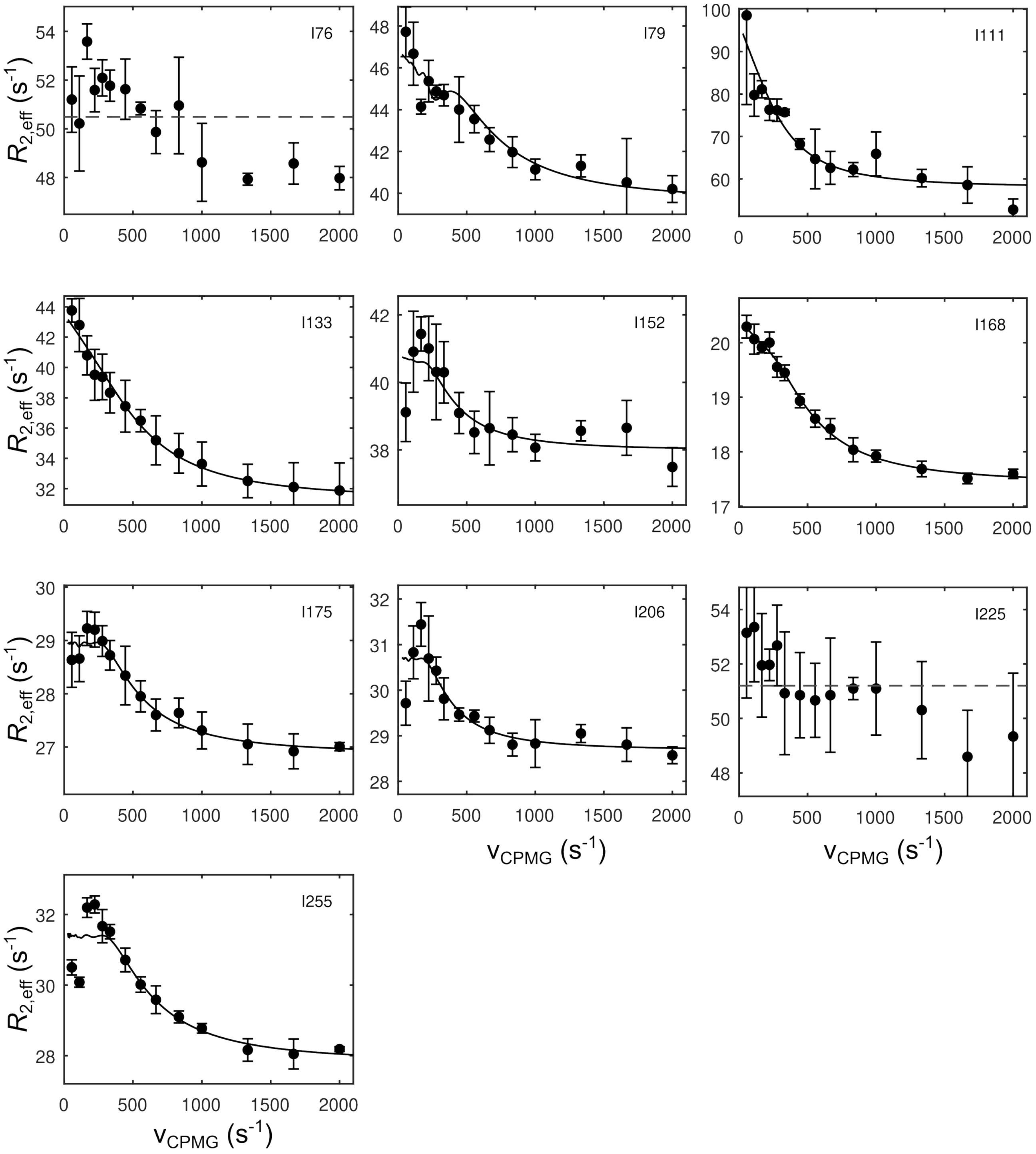


**Figure S4: MQ relaxation dispersion data for I-labeled Rrp45 bound to Rrp41.** The fits (solid lines) are based on a two-state model as described in the *Method* section. Reliable parameter determination is not expected due to scarcity of data. Dashed lines indicate absence of relaxation dispersion.

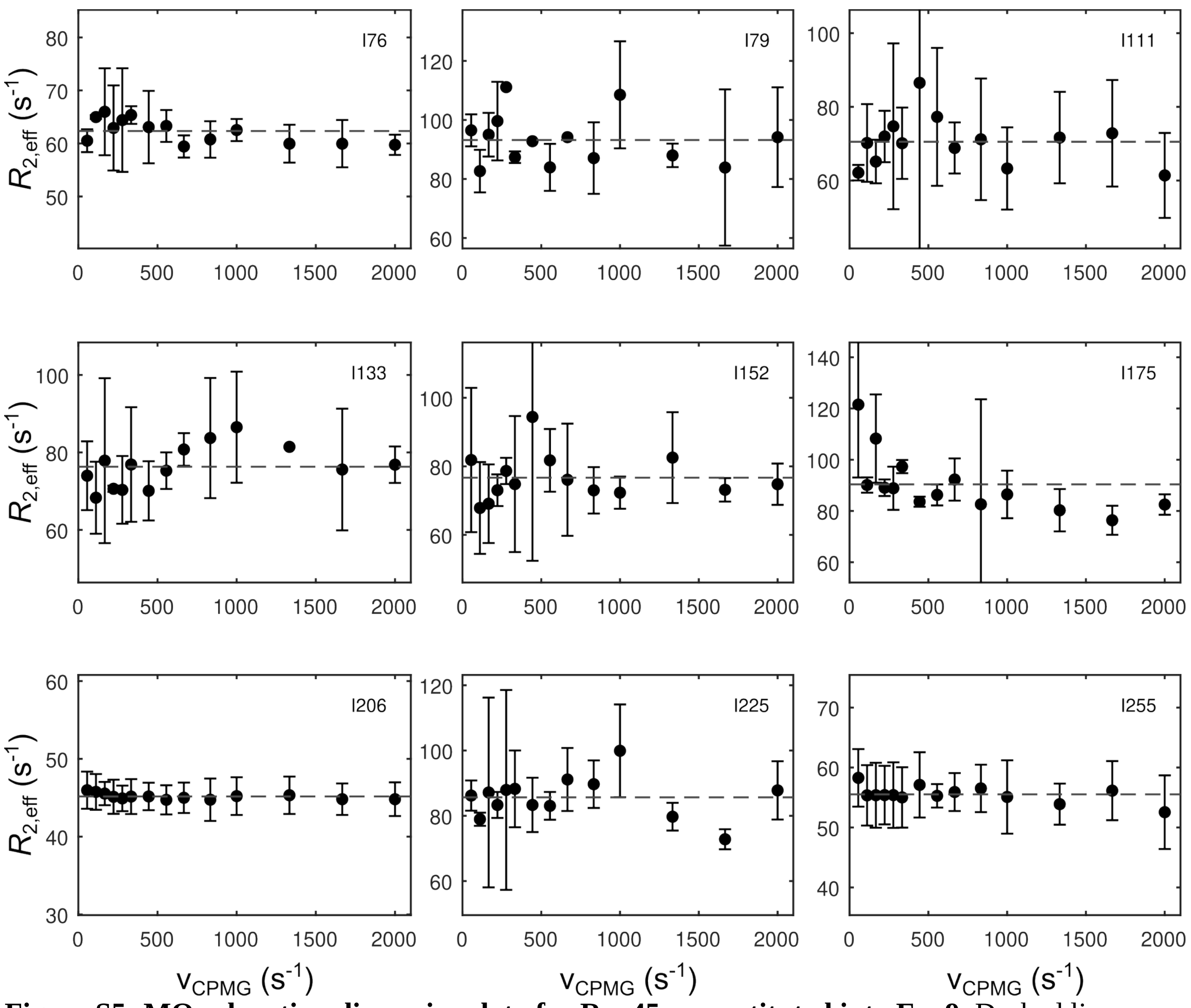


86 **Figure S5: MQ relaxation dispersion data for Rrp45 reconstituted into Exo9.** Dashed lines
87 indicate absence of relaxation dispersion.

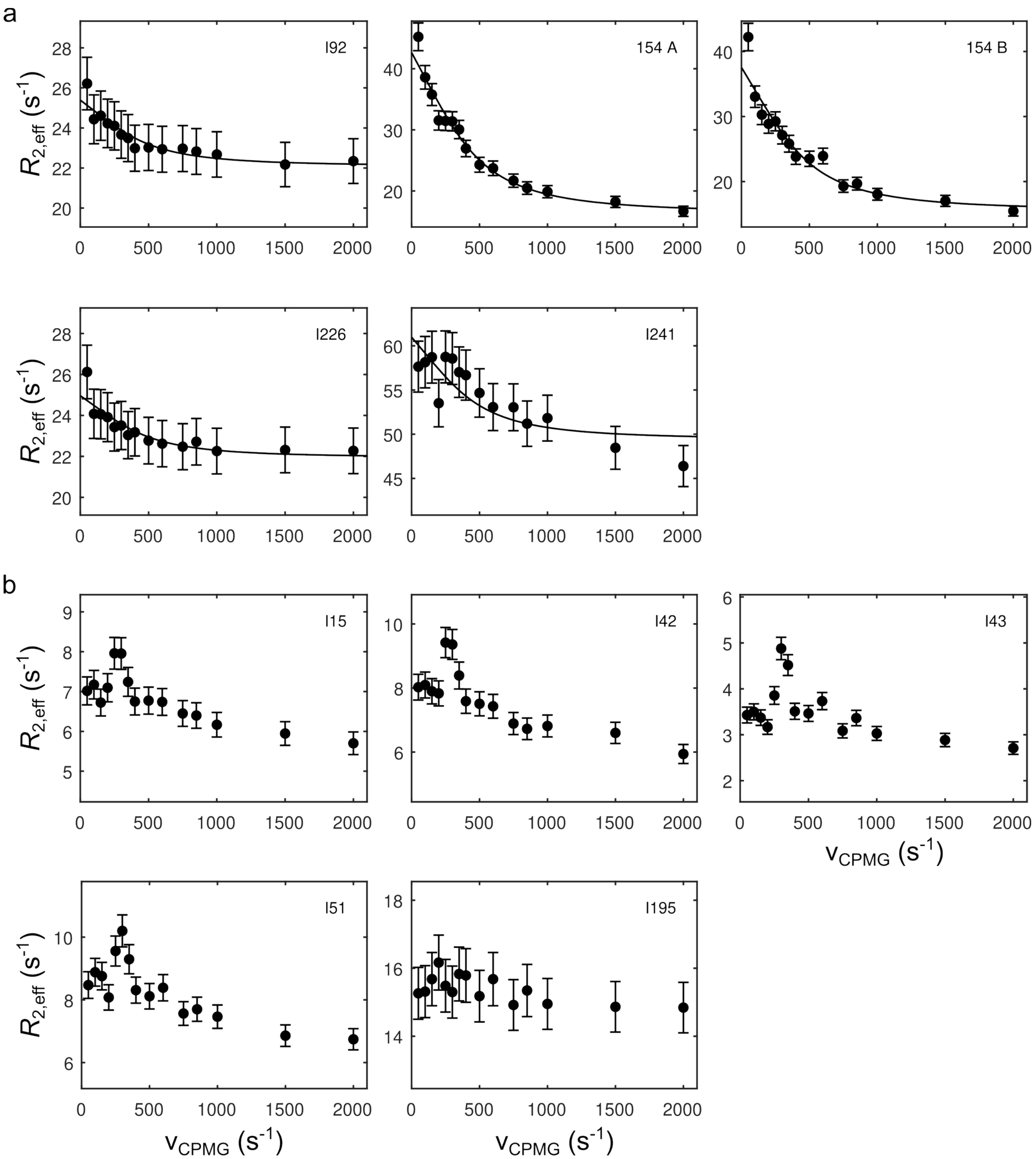


**Figure S6: Global fit for relaxation dispersion data of Rrp40. (a)** Resonances included in the global fit based on a two-state model as described in the *Method* section. **(b)** Resonances excluded from the global fit due to overlap or lack of relaxation dispersion. See **Table S7** for fitting parameters.

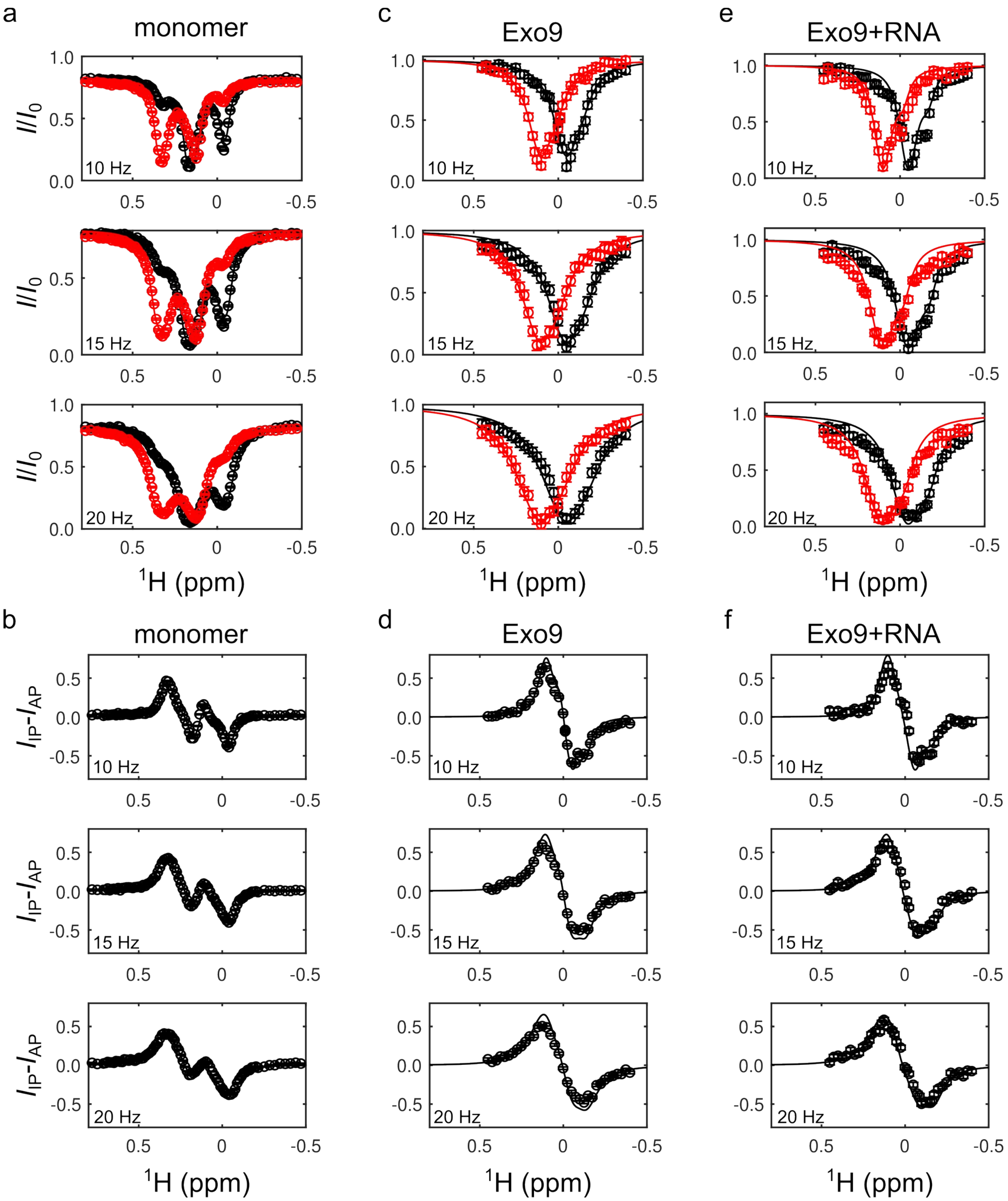


**Figure S7: $^{1}$H CEST profiles of Rrp40-I154 for IP magnetization (red) and AP magnetization (black) for (a) the monomer and when reconstituted into (c) Exo9 and (e) Exo9+RNA and difference profiles for (b) the monomer and when reconstituted into (d) Exo9 and (f) Exo9+RNA.** Data was acquired at $B_1$ fields of 10, 15 and 20 Hz. The fits are to a two-state exchange model as described in the method section. See **Table S8** for fitting parameters.

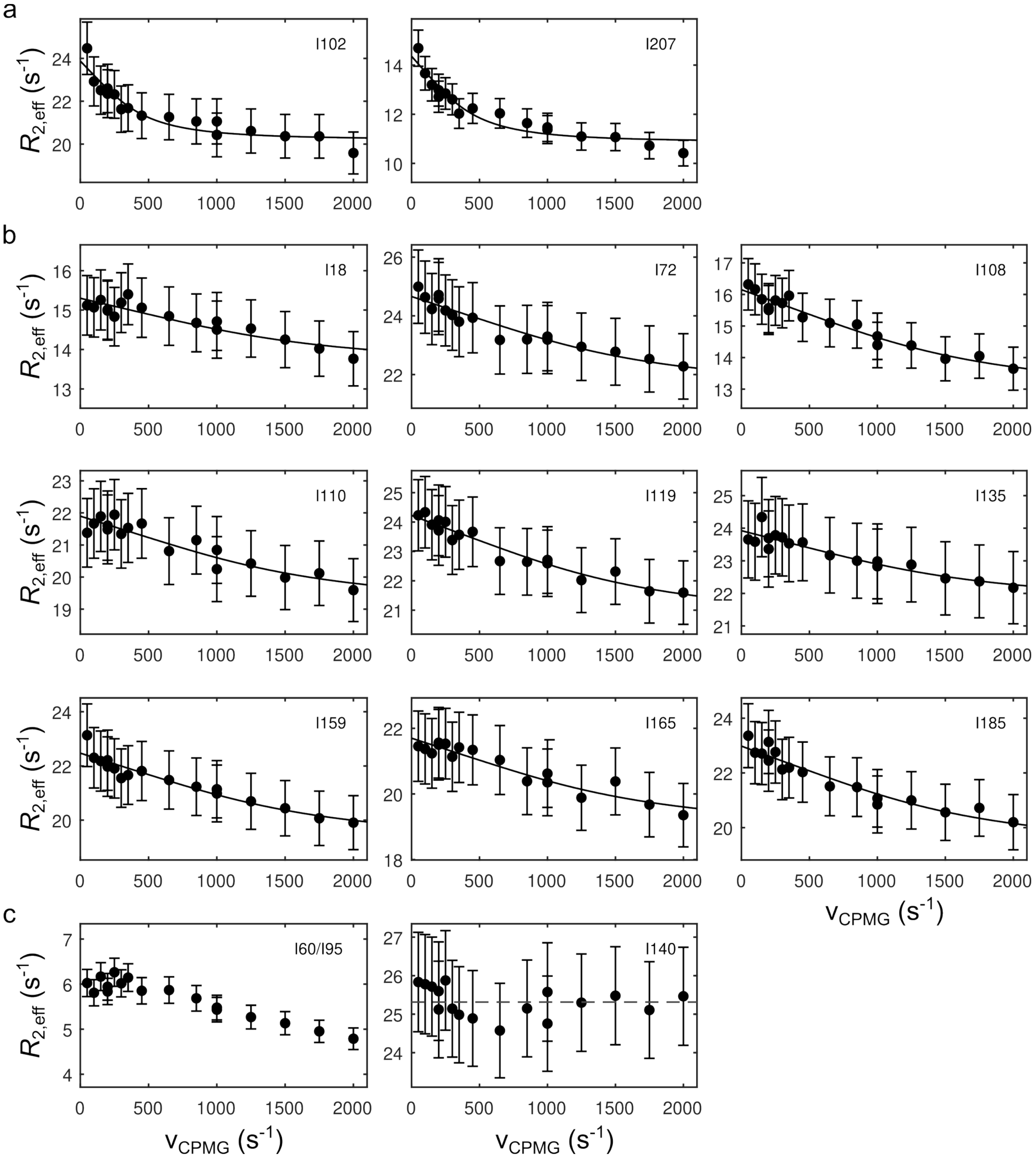


**Figure S8: Global fit for relaxation dispersion data of Csl4. (a)** Class A resonances and **(b)** class B resonances. The fits are based on a two-state model as described in the *Method* section. $k_{ex}$ and $p_B$ are global parameters for residues in the same class, $\Delta\nu_C$ and $R_{2,inf}$ are residues-specific parameters. **(c)** I60/I95 were not included in the fit due to peak overlap. I140 does not display any relaxation dispersion. See **Table S9** for fitting parameters.

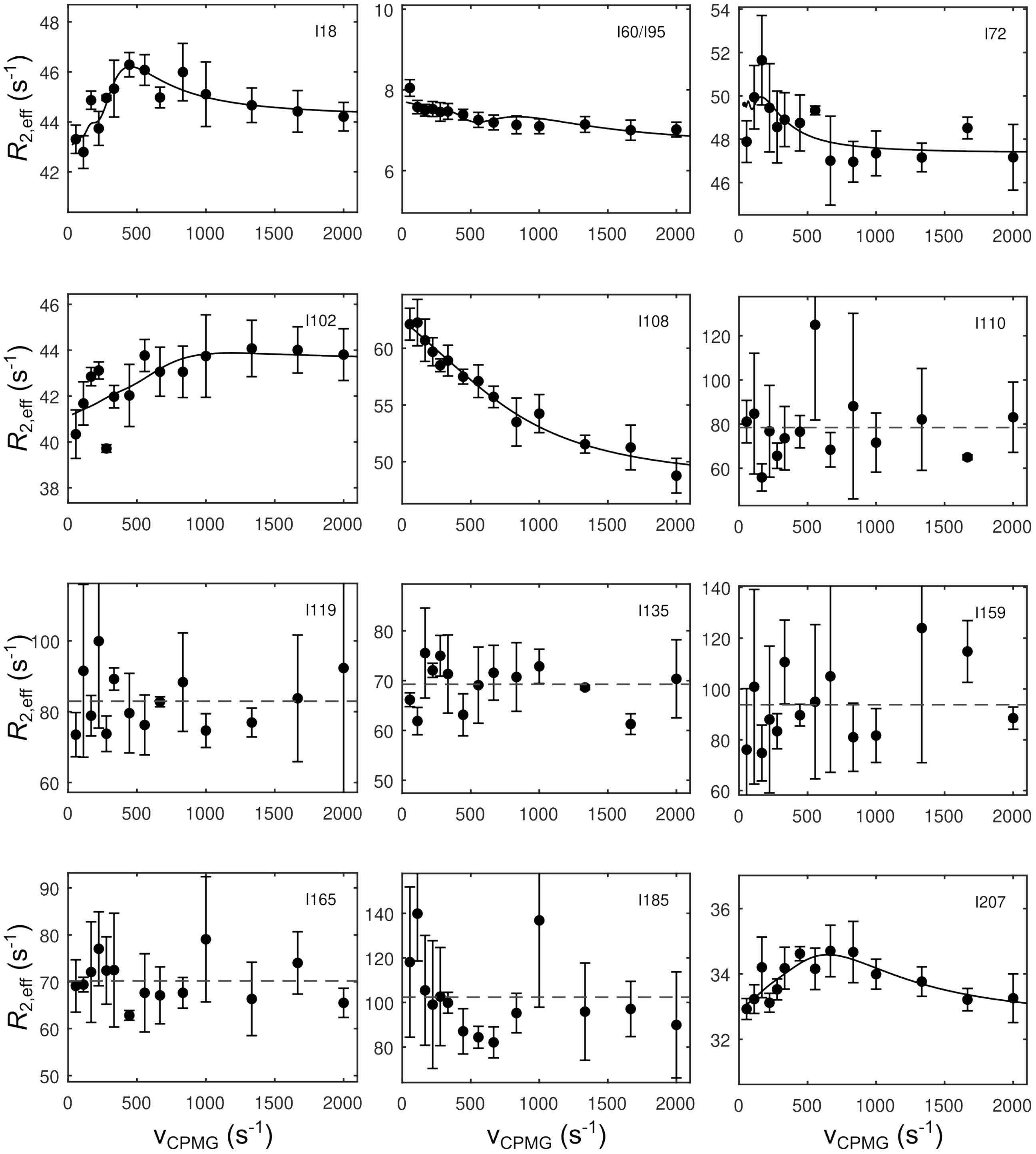


**Figure S9: MQ relaxation dispersion data for Csl4 reconstituted into Exo9.** The fits (solid lines) are based on a two-state model as described in the *Method* section. Reliable parameter determination is not expected due to scarcity of data. Dashed lines indicate absence of relaxation dispersion.

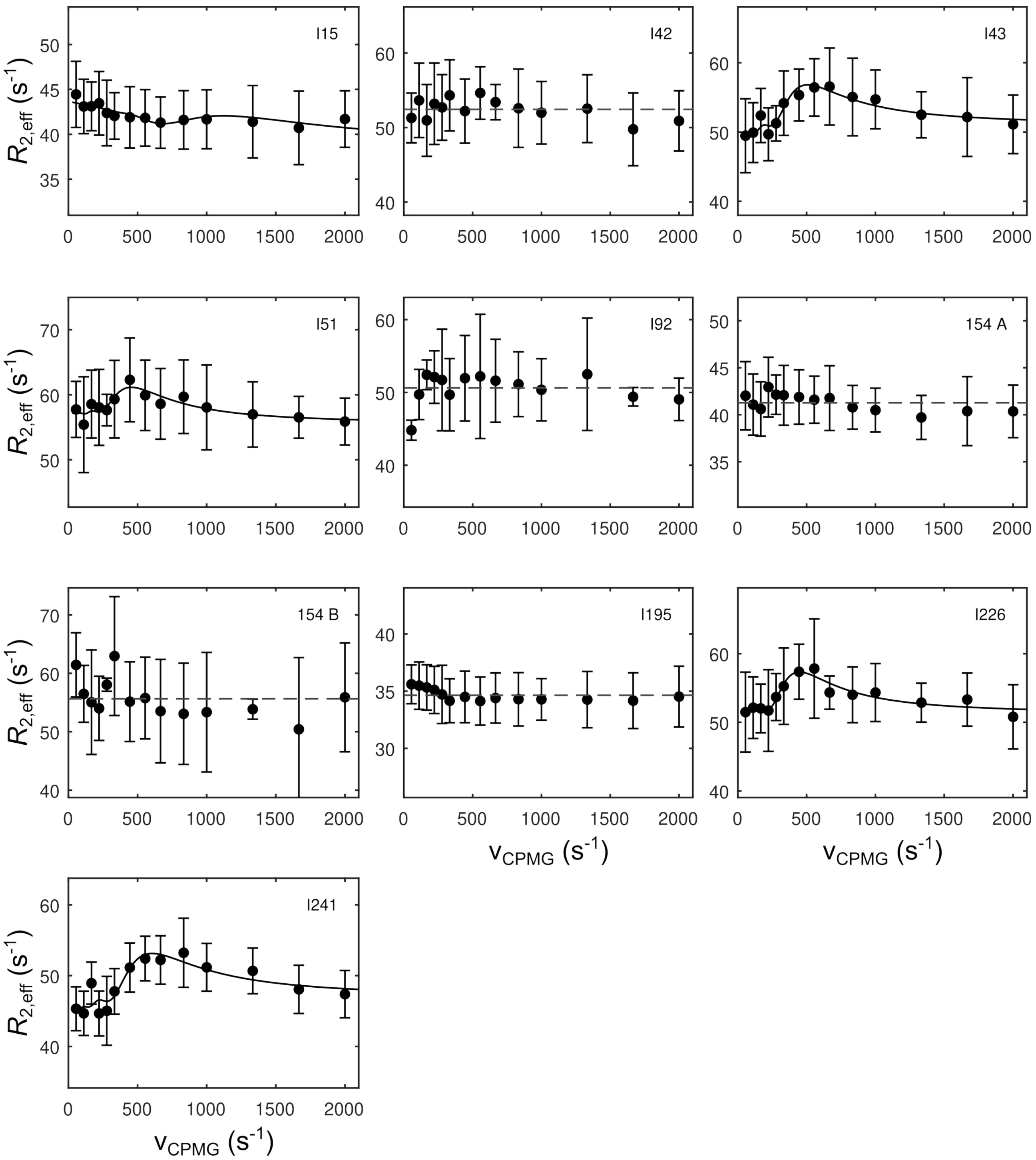


**Figure S10: MQ relaxation dispersion data for Rrp40 reconstituted into Exo9.** The fits (solid lines) are based on a two-state model as described in the *Method* section. Reliable parameter determination is not expected due to scarcity of data. Absence of dynamics are indicated by a dashed line.

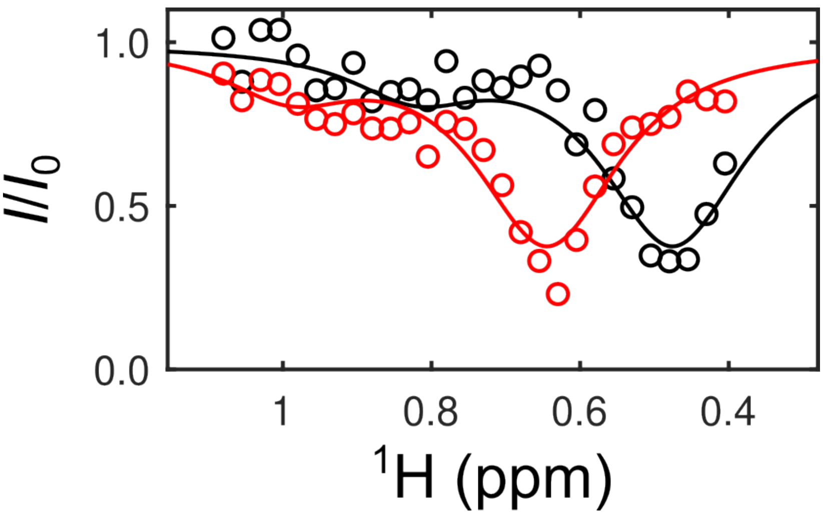


**Figure S11: $^1$H CEST of Rrp40-I51 in Exo9, acquired with $B_1$ = 10 Hz.** A broad exchange peak is observed at chemical shifts corresponding to monomer resonances indicating slow exchange between free and exosome-bound Rrp40.

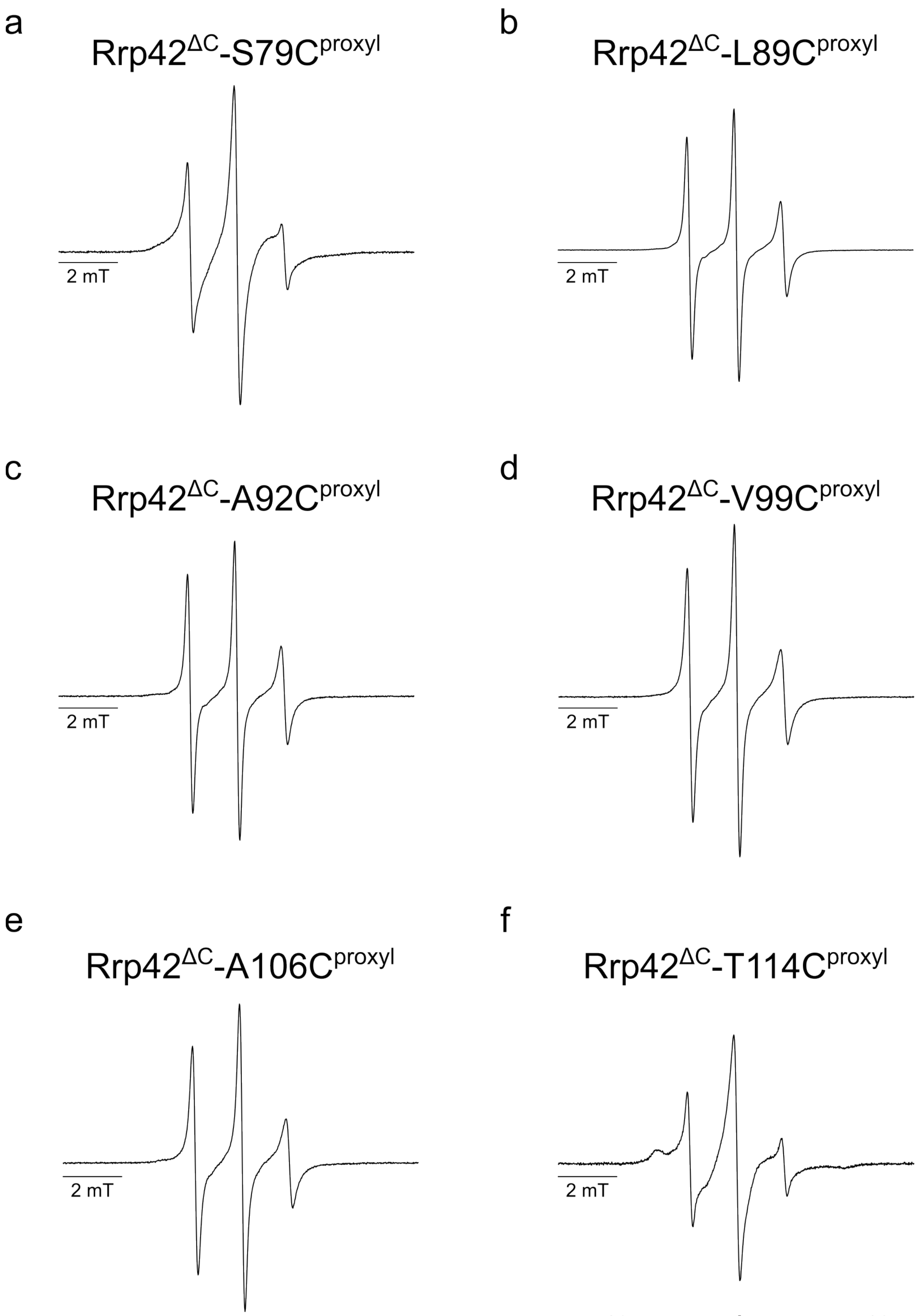


**Figure S12:** CW X-band EPR spectra of monomeric (a) Rrp42$^{\Delta C}$-S79C$^{proxyl}$, (b) Rrp42$^{\Delta C}$-L89C$^{proxyl}$ and (c) Rrp42$^{\Delta C}$-A92C$^{proxyl}$, (d) Rrp42$^{\Delta C}$-V99C$^{proxyl}$, (e) Rrp42$^{\Delta C}$-A106C$^{proxyl}$ and (f) Rrp42$^{\Delta C}$-T114C$^{proxyl}$ acquired at 294 K. The protein concentrations and labeling efficiencies are indicated in the **Table S1**.

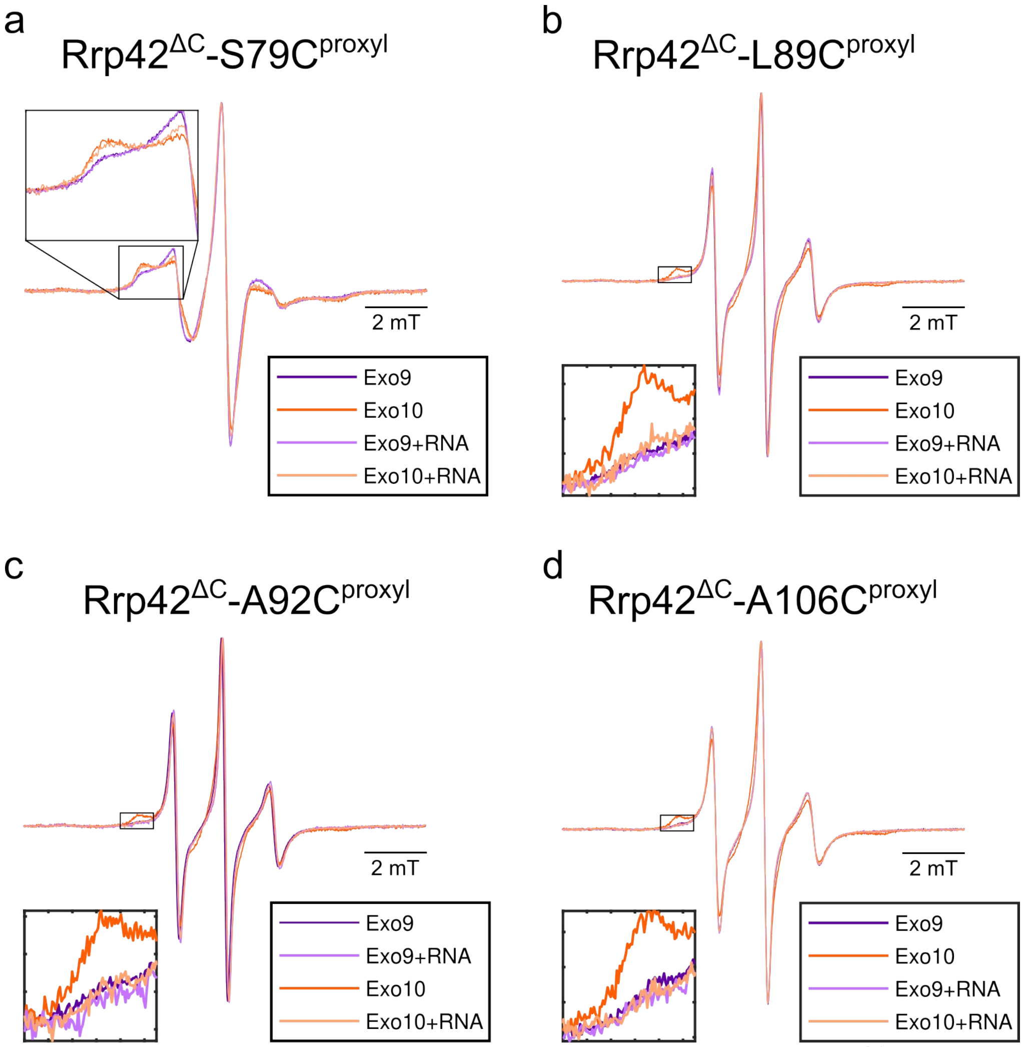


**Figure S13: Normalized CW X-band EPR spectra of spin-labeled Rrp42$^{\Delta C}$ in positions (a) S79C$^{proxyl}$**, located at the edge of Rrp42-EL and at **(a)** L89C$^{proxyl}$, **(b)** A92C$^{proxyl}$ and **(c)** A106C$^{proxyl}$, located in the center of Rrp42-EL acquired at 294 K, in Exo9 (purple) and Exo10 (orange) with (dark colors) and without (light colors) RNA. Insets: Zoom into the low field spectral region of the immobile fraction.

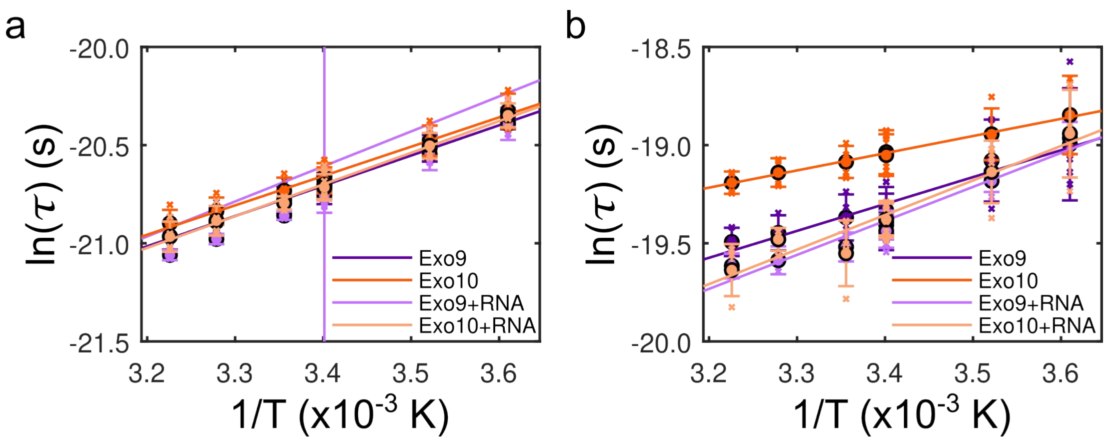


**Figure S14: Arrhenius plot** for the average populations of the **(a)** mobile and **(b)** immobile fractions of Rrp42-EL (positions V99C$^{proxyl}$, L89C$^{proxyl}$, A92C$^{proxyl}$ and A106C$^{proxyl}$) in Exo9 (purple) and Exo10 (orange) with (dark colors) and without (light colors) RNA. Circles are averages of measurements for the four different proxyl position in the loop (crosses). Error bars represent ±1 SD and fits are based on the Arrhenius equation. See **Table S14** for fitting parameters.
